# Structure and dynamics of Odinarchaeota tubulin and the implications for eukaryotic microtubule evolution

**DOI:** 10.1101/2021.10.22.465531

**Authors:** Caner Akıl, Samson Ali, Linh T. Tran, Jeremie Gaillard, Wenfei Li, Kenichi Hayashida, Mika Hirose, Takayuki Kato, Atsunori Oshima, Kosuke Fujishima, Laurent Blanchoin, Akihiro Narita, Robert C. Robinson

**Affiliations:** Research Institute for Interdisciplinary Science, Okayama University, Okayama 700-8530, Japan; Tokyo Institute of Technology, Earth-Life Science Institute (ELSI), Tokyo, 152-8551, Japan; Division of Biological Science, Graduate School of Science, Nagoya University, Furo-cho, Chikusa-ku, Nagoya, 464-8601, Japan; University of Grenoble-Alpes, CEA, CNRS, INRA, Interdisciplinary Research Institute of Grenoble, Laboratoire de Phyiologie Cellulaire & Végétale, CytoMorpho Lab, 38054 Grenoble, France; National Laboratory of Solid State Microstructure, Department of Physics, Collaborative Innovation Center of Advanced Microstructures, Nanjing University, 210093 Nanjing, China; Cellular and Structural Physiology Institute (CeSPI), Nagoya University, Furo-cho, Chikusa-ku, Nagoya 464-8601, Japan; Institute for Protein Research, Osaka University, Osaka, 565-0871, Japan; Department of Basic Medicinal Sciences, Graduate School of Pharmaceutical Sciences, Nagoya University, Furo-cho, Chikusa-ku, Nagoya 464-8601, Japan; Graduate School of Media and Governance, Keio University, Fujisawa, 252-0882, Japan; Université de Paris, INSERM, CEA, Institut de Recherche Saint Louis, U 976, CytoMorpho Lab, 75010 Paris, France; School of Biomolecular Science and Engineering (BSE), Vidyasirimedhi Institute of Science and Technology (VISTEC), Rayong, 21210, Thailand

## Abstract

Tubulins are critical for the internal organization of eukaryotic cells, and understanding their emergence is an important question in eukaryogenesis. Asgard archaea are the closest known prokaryotic relatives to eukaryotes. Here, we elucidated the apo and nucleotide-bound X-ray structures of an Asgard tubulin from hydrothermal-living Odinarchaeota (OdinTubulin). The GTP-bound structure resembles a microtubule protofilament, with GTP bound between subunits, coordinating the “+” end subunit through a network of water molecules and unexpectedly by two cations. A water molecule is located suitable for GTP hydrolysis. Time course crystallography and electron microscopy revealed conformational changes on GTP hydrolysis. OdinTubulin forms tubules at high temperatures, with short curved protofilaments coiling around the tubule circumference, more similar to FtsZ, rather than running parallel to its length, as in microtubules. Thus, OdinTubulin represents an evolution intermediate between prokaryotic FtsZ and eukaryotic microtubule-forming tubulins.

## INTRODUCTION

The tubulin/FtsZ/CetZ superfamily of proteins polymerize into filaments for which nucleotide-dependent dynamics and curvature are critical to their functions. The prokaryotic GTP-hydrolyzing FtsZ and CetZ form homo-filaments, which adopt straight and curved conformations (*1*–*4*). These filaments are part of the ring systems that constrict during prokaryotic cell division (*5*). The eukaryotic microtubule-forming tubulins have resulted from series of gene duplications (*6*), and have diverged significantly from FtsZ (*7*) and CetZ (*2*). The γ-tubulin ring complex patterns the microtubule (*8*), which typically comprises 13 parallel strands. Each straight strand (protofilament) nucleates from a single subunit of γ-tubulin via incorporation of the obligate α/β tubulin heterodimers (*9*). Thus, in the cell, the microtubule nucleation step is separated at microtubule organizing centers from other assembly and disassembly dynamics. α-tubulin contains a non-exchangeable, non-hydrolyzing GTP-binding site (N-site), whereas β-tubulin (E-site) and γ-tubulin contain exchangeable and hydrolyzing GTP-binding sites. The switch from straight to curved protofilaments at microtubule + ends, following GTP hydrolysis, results in catastrophe disassembly (*10, 11*).

The Asgard archaea superphylum have been proposed to be the closest prokaryotic relatives to eukaryotes (*12*). Their genomes, which were mainly obtained from metagenomic studies, include genes which have homology to eukaryotic signature protein (ESP) encoding genes. These ESPs were previously thought to be exclusive to eukaryotes, before the genomic characterization of the first Asgard archaea, Candidatus Lokiarchaeota (*13*). Thus, Asgard archaea genomes have become valuable resources to understand pre-eukaryotic protein machineries at the functional level, such as the ESPs studied in actin dynamics (*14*–*16*) and membrane fusion (*17*). The Candidatus Odinarchaeota archaeon LCB_4 (Odin) metagenome-assembled genome (MAG, GenBank accession number MDVT00000000.1) encodes two genes predicted to be FtsZ homologs (OLS17704.1 and OLS17546.1), and also possesses a single gene (OdinTubulin, OLS18786.1) that has greater homology to eukaryotic tubulin rather than to prokaryotic FtsZ (*12*). However, the properties of OdinTubulin are currently unknown.

## RESULTS

### X-ray structure of OdinTubulin

To address whether OdinTubulin is a genuine tubulin at the protein level, we expressed, purified, crystallized and determined the structure of OdinTubulin in the apo form, and bound to GTP or GDP (table S1 and fig. S1). Phylogenetic analysis, using structure alignment, confirmed that OdinTubulin has diverged significantly from FtsZ and CetZ, and branches in the same clade as eukaryotic tubulins (Fig. 1A) (*12*). Structure homology searches revealed that the GTP-bound OdinTubulin, refined at 1.62 Å (PDB 7EVB, capital letters refer to structures determined in this study) is most similar to α- and β-tubulins within a microtubule, regardless of the nucleotide state within the microtubule, rather than to non-polymerized tubulin subunits, or to FtsZ or CetZ (Fig. 1B, and table S2). OdinTubulin shares ∼35% sequence identity with the human α- and β-tubulins. Within the crystal packing, OdinTubulin subunits are arranged as in a microtubule protofilament (fig. S1 and S2). Superimposition of the OdinTubulin lower subunit (-) onto a eukaryotic microtubule GDP-containing β-tubulin subunit (*18*) revealed that the position of the OdinTubulin upper subunit (+) aligned closely with the proximal α-tubulin subunit from the microtubule protofilament (Fig. 1C) (*18*). By contrast, superimposition onto the microtubule structure containing a GTP mimetic in the β-tubulin subunit (*11*), or onto the β-tubulin subunit from the stathmin-bound curved protofilaments (*19*), aligned the α-tubulin subunits poorly to the OdinTubulin upper subunit (fig. S3, table S3 and movie S1 to S3).

**Fig. 1.**
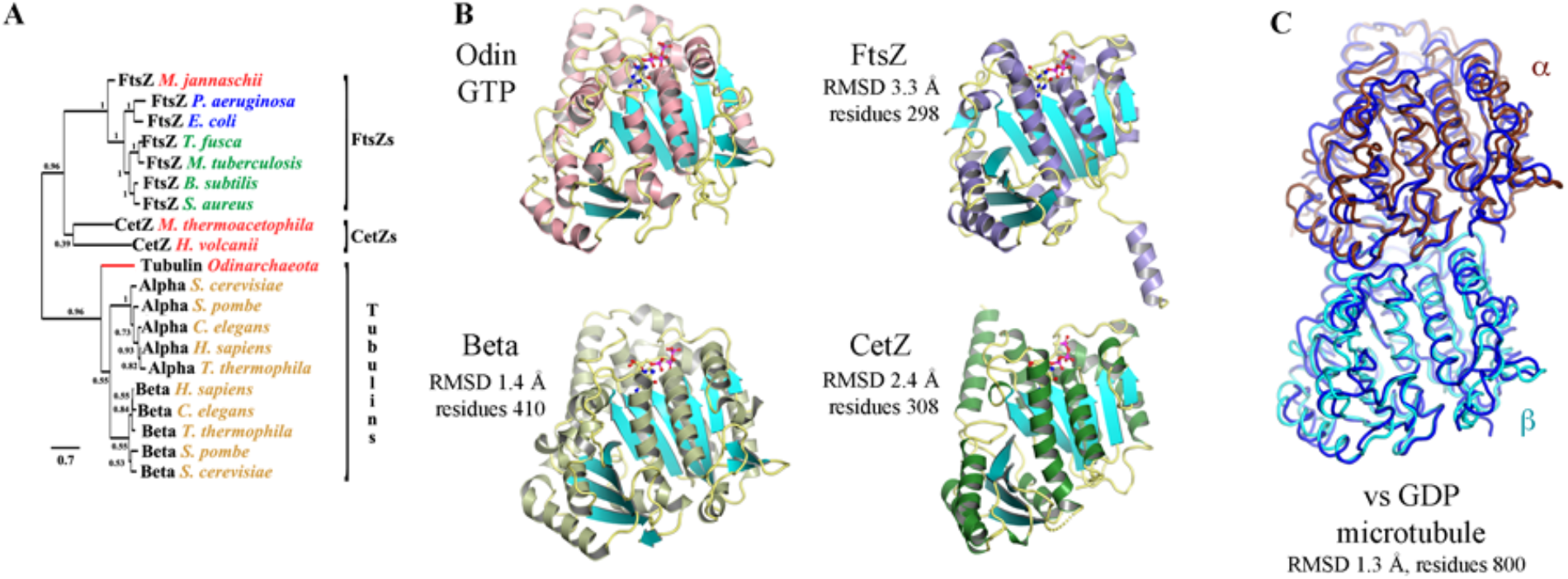
The crystal structure of OdinTubulin. (A) Phylogenetic analysis of OdinTubulin from structure-based sequence alignment in comparison to the prokaryotic cell division proteins, FtsZ and CetZ, and the eukaryotic microtubule-forming tubulins. (B) Comparison of the protomer structures of GTP-bound OdinTubulin (PDB 7EVB) to β-tubulin (PDB 6o2r) (*18*), CetZ (PDB 4b45) (*2*) and FtsZ (PDB 1w5a) (*4*). The matching numbers of residues and RMSD values indicate the relative structural similarities to OdinTubulin. (C) Superimposition of the two GTP-bound OdinTubulin symmetry-related subunits from the crystal packing (dark blue) onto two subunits of eukaryotic tubulin from the GDP-bound microtubule (PDB 6o2r) (*18*).

Similar to eukaryotic tubulin, OdinTubulin comprises an N-terminal domain (residues 1-202), and intermediate domain (residues 203-367) and a C-terminal domain (residues 368-424, Fig. 2A), as described for eukaryotic tubulin (*9*). The α7 helix and its preceding loop (blue) and the α8 helix and its preceding loop (red), which we term the “nucleotide sensor motif”, lies in the intermediate domain, connecting the nucleotide from the lower subunit (-) to the nucleotide in the upper subunit (+) (Fig. 2A and movie S4). In eukaryotes, the α7 helix is known to undergo a translation movement in response to the presence of different nucleotides and protofilament curvature (*20*). Taken together, these data indicate that the GTP-bound OdinTubulin crystals contain a straight microtubule-like tubulin protofilament stabilized by native nucleotide binding.

**Fig. 2.**
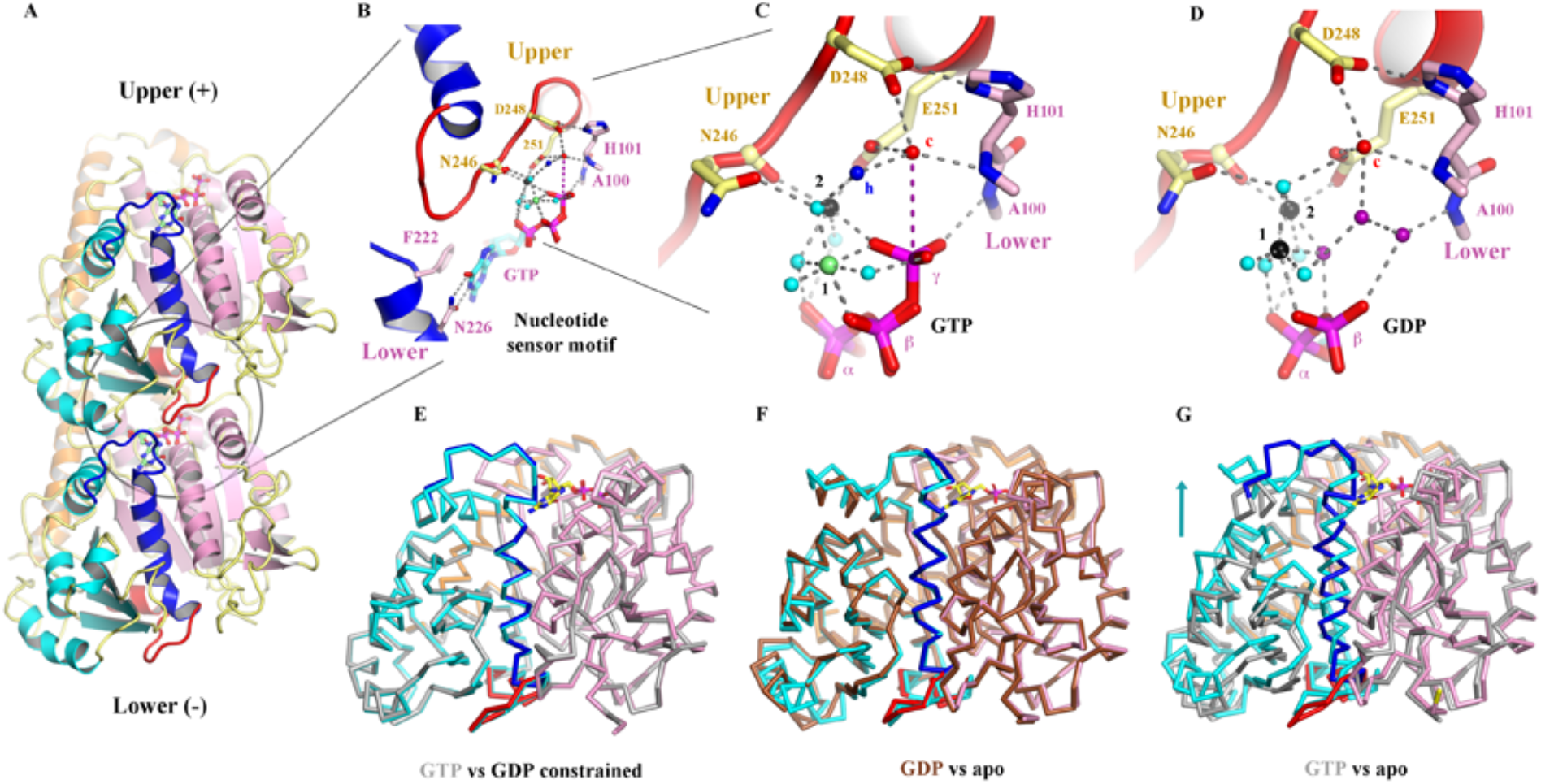
Structural implications for GTP hydrolysis. (A) The OdinTubulin protofilament in the crystal packing (PDB 7EVB). Two subunits of OdinTubulin are depicted. The α7 helix and preceding loop (blue) and α8 helix and preceding loop (red) comprise the nucleotide sensor motif, which connect the upper and lower GTP-binding sites (sticks). Secondary structure elements are colored by domain: N-terminal (pink), intermediate (cyan), and C-terminal (orange). The nucleotide sensor motif lies within the intermediate domain. See movie S4. (B) Enlargement of the GTP interactions. Only part of each nucleotide sensor motif is shown for clarity. Selected residues from the upper and lower subunits are labelled in yellow and pink, respectively. (C) Enlargement of the interactions around the GTP γ-phosphate. Black, lime green and cyan spheres indicate Na^+^, Mg^2+^ (numbered in black) and water molecules, respectively. The proposed hydrolytic water is shown as a red sphere and labeled “c”, and water molecule suitably placed to receive the hydrogen ion from the hydrolytic water is labeled “h” in blue. The purple dashed line indicates the route for nucleophilic attack on the GTP γ-phosphate. See movie S5. (D) The same region from the GDP-bound structure (PDB 7EVE). Three water molecules (purple) replace the GTP γ-phosphate, and both cations are assigned as Na^+^ based on bond distances and crystallization condition. See movie S5. (E-G) Superimposition of protomer structures. (E) GTP-bound OdinTubulin (grey, PDB 7EVB) overlaid on the constrained GDP-bound structure (colored, PDB 7EVE). (F) The unconstrained GDP-bound OdinTubulin (grey, PDB 7EVB) overlaid on the apo structure (colored, PDB 7EVG). (G) GTP-bound OdinTubulin (grey, PDB 7EVB) overlaid on the apo structure (colored, PDB 7EVG). The arrow highlights the conformational change for the intermediate domain. See movie S7.

### GTP binding

Inspection of the nucleotide-binding site revealed that GTP is bound to the “E” site, with the guanine moiety interacting with the sidechains of Phe222 and Asn226 from the N-terminal region of the nucleotide sensor motif, and the gamma phosphate is bound to the mainchain amide nitrogen from Ala100 from the lower subunit (Fig. 2B). The C-terminal region of the nucleotide sensor motif, from the upper subunit, interacts indirectly with the GTP phosphate groups through a bonding network of cations and water molecules (Fig. 2, B and C). There are two GTP-bound cations. The commonly observed ion that bridges the beta and gamma phosphates (“1” in Fig. 2C) and a second ion that directly coordinates Asn246, Glu251 and the GTP gamma phosphate (“2” in Fig. 2C). Cation 2-stabilized Glu251 orders a water molecule (“c” in Fig. 2, B and C), which is also in bonding distance of Asp248 from the upper subunit, and to the mainchain amide nitrogen from His101 in the lower subunit. Water “c” is 4.1 Å from the GTP gamma phosphorous atom, suitably positioned for straight-line nucleophilic attack for hydrolysis (movie S5). The cation binding sites (1 and 2) are promiscuous but, in the crystal structures, preferentially bind Mg^2+^ and K^+^/Na^+^, respectively (fig. S4).

### Proposed hydrolysis mechanism

The GTP-binding site arrangement indicates a probable hydrolysis mechanism, whereby and Asp248 and/or Glu251 activate the hydrolytic water “c”. The activated water will be directed by the mainchain amide nitrogen from His101 and the sidechain of Glu251, which may swivel while remaining bound to cation 2, resulting in nucleophilic attack on the gamma phosphorous atom, leading to hydrolysis (fig. S5). Another water molecule “h” is suitably placed to receive the hydrogen ion from the hydrolytic water (blue, Fig. 2C and movie S5). A crystal structure containing 100% GDP in the nucleotide-binding site (PDB 7EVE, refined at 2.0 Å) revealed that the phosphate ion is released following hydrolysis, and is replaced by three water molecules and the hydrolytic water binding site is occupied (Fig. 2D and movie S5), without eliciting significant conformational changes to the OdinTubulin protomer or protofilament structures (Fig, 2E and fig. S1). We propose that the exchange of the covalently bound γ-phosphate for three water molecules, following hydrolysis, results in weakening the binding between the upper and lower OdinTubulin subunits in this region, producing strain in the protofilament.

### Conformational changes on GTP hydrolysis

Incubation of GTP-soaked crystals in the presence of Na^+^ over time resulted in an initial increase in the bound GDP:GTP ratio (3 days, Fig. 3) followed by a blurring of the electron density for the intermediate domain (2 months), indicating a slow structural transition within the crystals. A single GTP-soaked OdinTubulin crystal (5 mM GTP, 1 mM MgCl_2_ and 0.1 M KCl, 1h) was frozen and confirmed to contain a ratio of GTP:GDP of ∼9:1 by X-ray crystallography. Subsequently, the crystal was thawed and re-equilibrated in crystallization conditions supplemented by 1 mM MgCl_2_, 0.1 M KCl, and 0.2 mM sodium acetate in the absence of nucleotide (2 weeks at 20 °C), before being refrozen and the crystal structure determined (PDB 7F1B, refined at 2.40 Å). The resulting 100% GDP-bound conformation represents a second class of OdinTubulin structure which is similar to structures we also determined in the apo state (PDB 7EVG, refined at 2.48 Å), or partially bound to background GDP that resulted from the purification protocol (∼60%, PDB 7EVH, refined at 2.50 Å; Fig. 2F, fig. S1 and S6, and movie S6). We interpret this alternate OdinTubulin conformation to be that after the structural transitions resulting from GTP hydrolysis to GDP, allowing for release of the nucleotide.

**Fig. 3.**
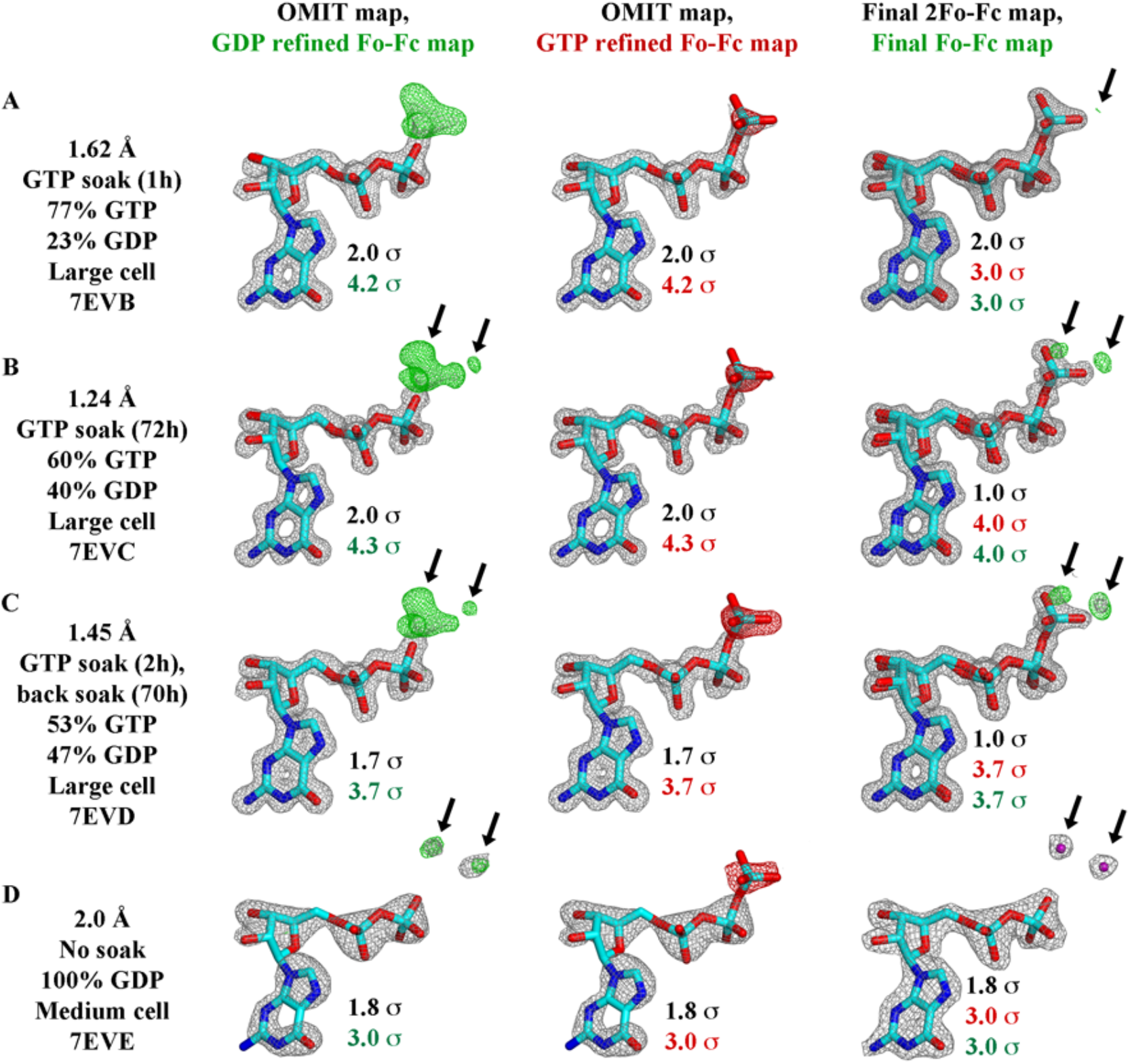
GTP hydrolysis followed by X-ray crystallography. Structures determined (A) 1 h or (B) 72 h after soaking with 10 mM GTP showed a decrease in the bound GTP:GDP ratio. (C) Back soaking the crystals for 70 h, decreased the ratio further. (D) The structure of a non-soaked crystal with 100% GDP bound to OdinTubulin arranged in the regular protofilament packing, similar to the GDP-bound subunits within a microtubule, which is stabilized by a different crystal unit cell (fig. S1 and S2). The maps are contoured at levels indicated by the color-coded sigma levels. Left column, the structures were refined with GDP in the nucleotide-binding site. Middle column, the structures were refined with GTP in the nucleotide-binding site. Left column, the structures were refined with final GTP/GDP ratios in the nucleotide-binding sites. Green (+) and red (-) Fo-Fc density indicate the need for more or less atoms, respectively. The arrows indicate the position of two ordered water molecules that appears following γ-phosphate release after hydrolysis, which are shown in purple (D, right). The third ordered water molecule (Fig. 2D), which appears after hydrolysis, is bound to the metal ions and occupies a similar position to an oxygen from the GTP γ-phosphate. Thus, this water does not appear in the difference maps. This water has weaker electron density compared to the two waters detailed in this figure, and likely partial occupancy (movie S5).

Comparison of apo/GDP-bound structure with the GTP-bound structure, revealed that the intermediate domain, including the nucleotide sensor motif, moves relative to the N-terminal and C-terminal domains (Fig. 2G and movie S7). This conformational change alters the interactions between protomer subunits and likely results in a curving of the protofilament, unless the straight form is stabilized by inter-protofilament interactions as observed for the central portion of the microtubule, or in the OdinTubulin crystal packing (fig. S2). By contrast the 100% GDP bound OdinTubulin (PDB 7EVE) adopts a GTP-like conformation stabilized by a different crystal packing (fig. S1 and S2). We interpret the 7EVE structure to be the GDP-bound strained structure prior to the conformational change. The role of the nucleotide sensor motif in allosterically linking the occupancy of the nucleotide-binding site between adjacent protomers is likely twofold: firstly, in ensuring GTP-bound monomers are preferentially added to a growing filament, and secondly, in cooperatively coordinating the conformational change throughout a protofilament following hydrolysis and phosphate release.

### Conservation with microtubule-forming tubulins

Since, the proposed hydrolytic and ion-binding residues from the nucleotide sensor motif are conserved between OdinTubulin and α-tubulin (Fig. 4A), we propose that GTP hydrolysis in microtubules may proceed via the same mechanism involving two cations and a strained intermediate where the phosphate ion is released following hydrolysis and replaced by three water molecules (fig. S5). Support for this mechanism can be found in the structures of eukaryotic tubulins. A water molecule is found bound to α-tubulin Glu254 in the sequestered α/β-tubulin dimer (*21*), equivalent to the hydrolytic water bound to Glu251 in OdinTubulin (compare Fig. 4B and 4C). α-tubulin Glu254 has been predicted to be involved in catalysis by comparison to the structure of FtsZ (*22*). The mainchain amide nitrogens of Ala99 and Gly100 (β-tubulin) adopt similar positions to Ala100 and His101 from the lower OdinTubulin subunit, which interact with the GTP γ-phosphate and water “c”, respectively (compare Fig. 2B and 2D).

**Fig. 4.**
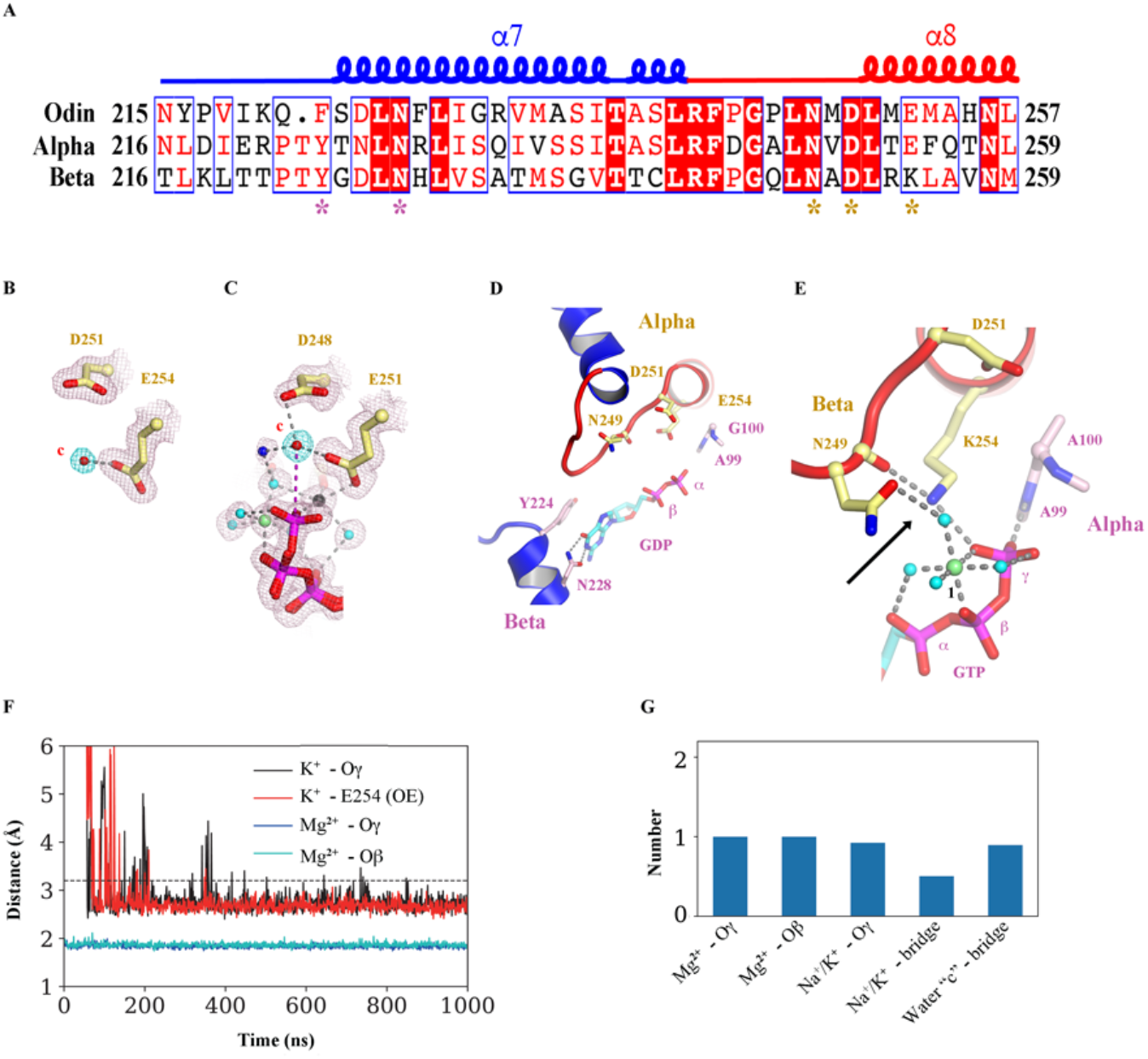
Similarities in OdinTubulin and microtubule nucleotide interactions. (A) Conservation in the sequence of the nucleotide sensor motif from OdinTubulin and human α- and β-tubulins. Colored stars below the alignment indicate the residues highlighted in Fig. 2, B to D and this Fig. (B to E). (B) A water molecule is found bound to α-tubulin Glu254 in the sequestered α/β-tubulin dimer (PDB 6s8k), equivalent to (C) the hydrolytic water “c” bound to Glu251 in OdinTubulin. The 2Fo-Fc electron density maps contoured at 1 σ (pink) and the density around potential hydrolytic water (cyan). (D) Subunits within a GDP-bound microtubule (PDB 6o2r) in a similar conformation to Fig. 2B showing structural similarity. (E) β-tubulin interactions with bound-GTP α-tubulin (PDB 6s8k) in a similar orientation to Fig. 2C. The cation-bound, hydrolysis-guiding residue Glu251 from OdinTubulin is substituted by a basic residue (Lys254, indicated by the arrow) in β-tubulin. (F) Coordination of metal ions in molecular dynamics simulations at the GTP exchangeable site of a microtubule for a representative. A magnesium ion is stably coordinated via oxygen atoms from GTP β (cyan) and γ (blue) phosphates throughout the 1 μs simulation. K^+^ becomes associated with an oxygen atom from the GTP γ -phosphate (black) and Glu254 (red). Similar results were obtained when Na^+^ became coordinated at the same site. (G) Occupancies of the metal ions at each site during the simulation. Stable occupancies of the GTP-bound Mg^2+^ and K^+^ or Na^+^ were observed. Approximately half the time the K^+^ or Na^+^ were jointly coordinated by the GTP γ-phosphate and Glu254 (Na^+^/K^+^ bridge). Finally, a water molecule(s) was located at position “c” between Glu254 and the phosphorous atom of the GTP γ -phosphate (water “c” – bridge).

Furthermore, in the 3.3 Å cryoEM structure of GDP-bound E-site microtubule (*18*) adopts a similar conformation around the nucleotide (compare Fig. 2B and 4D). However, identification of low molecular weight species, such as water molecules, has not been possible at the resolution of the microtubule EM maps. In the sequestered α/β-tubulin dimer (*21*), the hydrolysis-activating residue Glu251 from OdinTubulin is substituted by a basic residue (Lys254) in β-tubulin, which places a positive charge in the same location as magnesium ion 2 (compare Fig. 2C and 4E). This results in a lack of hydrolytic activity by the β-tubulin in the N-site, but also indicates that a positive charge is acceptable, in eukaryotic tubulins, in bridging the carbonyl of Asn249 (Asn246 in OdinTubulin, Fig 2C) and the GTP gamma phosphate (Fig. 4E).

To add weight to the prediction that there is a common mechanism of hydrolysis between OdinTubulin and microtubules, we carried out molecular dynamics (MD) simulations on restrained α/β-tubulin subunits at the “E” site interface in a background of magnesium, sodium, and potassium ions. In the 1 μs simulation, a magnesium ion stably associated with the oxygen atoms from the β and γ GTP phosphates (Fig. 4F), similarly to the OdinTubulin crystal structure (Fig. 2C). Furthermore, a potassium or sodium ion quickly became within bonding distance of oxygen atom of the GTP γ-phosphate and a carboxyl atom of Glu254 (Fig. 4F), similar to cation 2 in the OdinTubulin structure (Fig. 2C). We measured the occupancy of the ions at these sites during the simulation (Fig. 4G). The Mg^2+^ and K^+^/Na^+^ binding sites, in interacting with GTP, were fully occupied during the simulation. Approximately, 40% of the time a K^+^/Na^+^ occupied the bridging site between oxygen atom of the GTP γ-phosphate and a carboxyl atom of Glu254 (Fig. 4G). Finally, a water molecule was observed at 100% occupancy in bridging Glu254 (Glu251 in OdinTubulin) and the phosphorous atom of the GTP γ-phosphate (water “c” – bridge, Fig. 4G and 2C). Taken together, the conservation in the important GTP-hydrolysis residues, which bind cations and the proposed hydrolytic water, combined with the occupancy of these sites in MD simulations, reinforce the hypothesis that GTP hydrolysis proceeds through similar mechanism in OdinTubulin and microtubules that involves two cations.

### OdinTubulin filament assembly

We observed the dynamics of OdinTubulin polymerization by interference reflection microscopy (IRM) (*23*). Unlike the sparse, straight, smoothly elongating microtubules (Fig. 5A and movie S8 and S9), OdinTubulin tended to form wider bundles of filaments, under similar conditions (Fig. 5B and movie S8 and S9). The elongation proceeded with a high rate of nucleation via polymerization and filament annealing. Lower concentrations of OdinTubulin led to shorter more uniform filaments, which could be assembled under a variety of cation conditions (Fig. 5, C to F and movie S8 and S9).

**Fig. 5.**
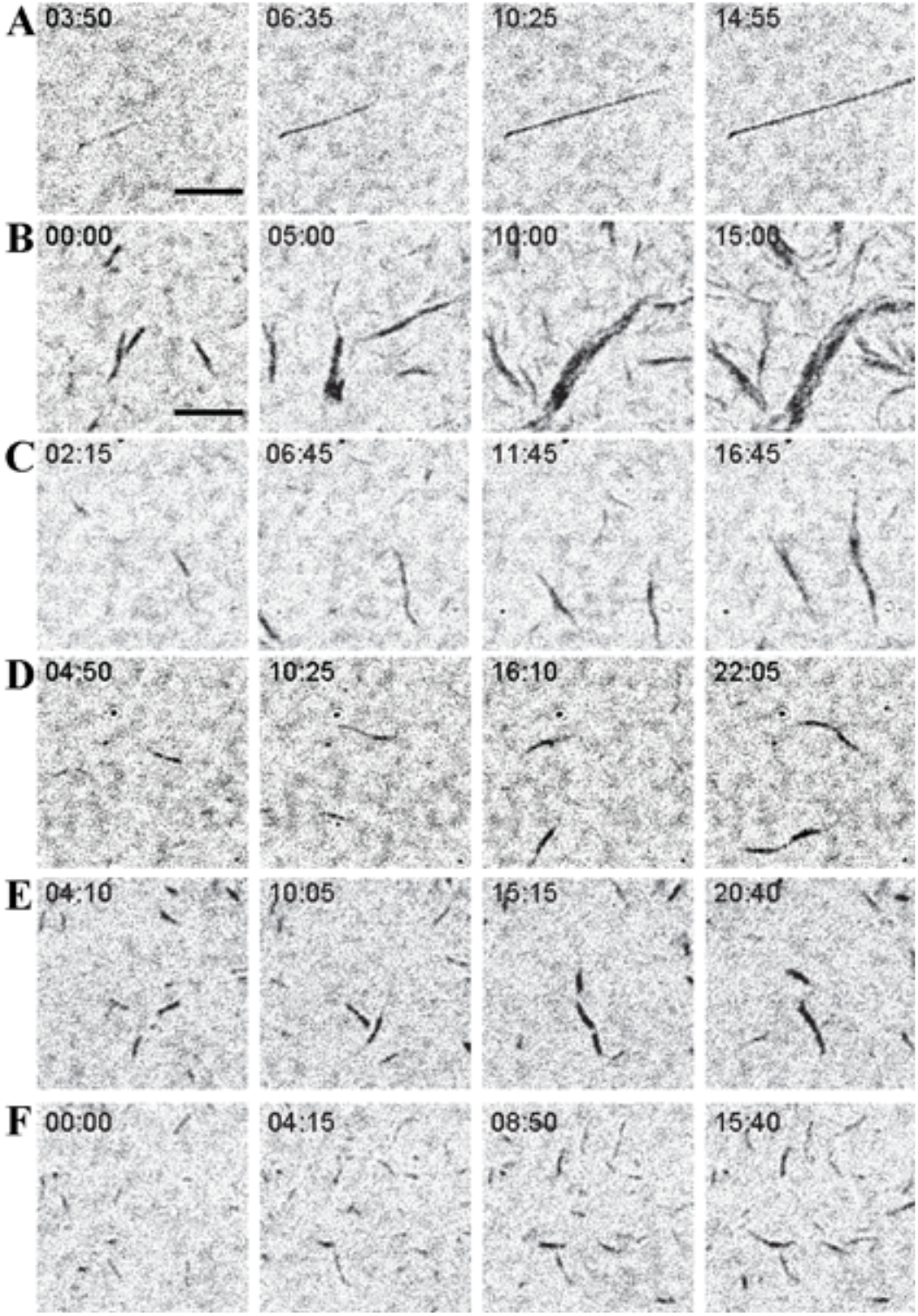
Polymerization of OdinTubulin followed by IRM. (A) Elongation of tubulin (15 μM) into microtubules in 100 mM Pipes-K, pH 6.9, 0.5 mM MgSO_4_, 0.5 mM EGTA, 10% glycerol, 0.7 mM GTP. (B-D) Polymerization of OdinTubulin at 6 μM, 2 μM and 1 μM, respectively, under the same solution conditions. (E) Polymerization of OdinTubulin (0.5 μM) in 100 mM Pipes-K, pH 6.9, 100 mM NaCl, 0.5 mM EGTA, 0.7 mM GTP or (F) supplemented with 0.5 mM MgSO_4_.

Observation of negatively stained polymers by electron microscopy (EM) indicated that two forms of polymer could be assembled. In the absence of Mg^2+^, bundles of straight protofilaments assembled in solutions containing the monovalent cations, K^+^ or Na^+^ (Fig. 6A). Inclusion of Mg^2+^ with a high concentration of Na^+^led to a mixture of bundled protofilaments and tubules (Fig. 6A). By contrast, Mg^2+^/K^+^ solutions, in the absence of Na^+^, produced exclusively tubules (Fig. 6B). Since, K^+^/Na^+^ and Mg^2+^ are able to bind to the GTP in OdinTubulin protofilaments (fig. S4), but hydrolysis is slow in the presence of Na^+^ and absence of Mg^2+^ (Fig. 3), we interpret the bundles to form from polymerized OdinTubulin protofilaments before significant GTP hydrolysis, whereas the tubules likely are formed simultaneously with GTP hydrolysis, allowing a structural transition to a curved morphology.

**Fig. 6.**
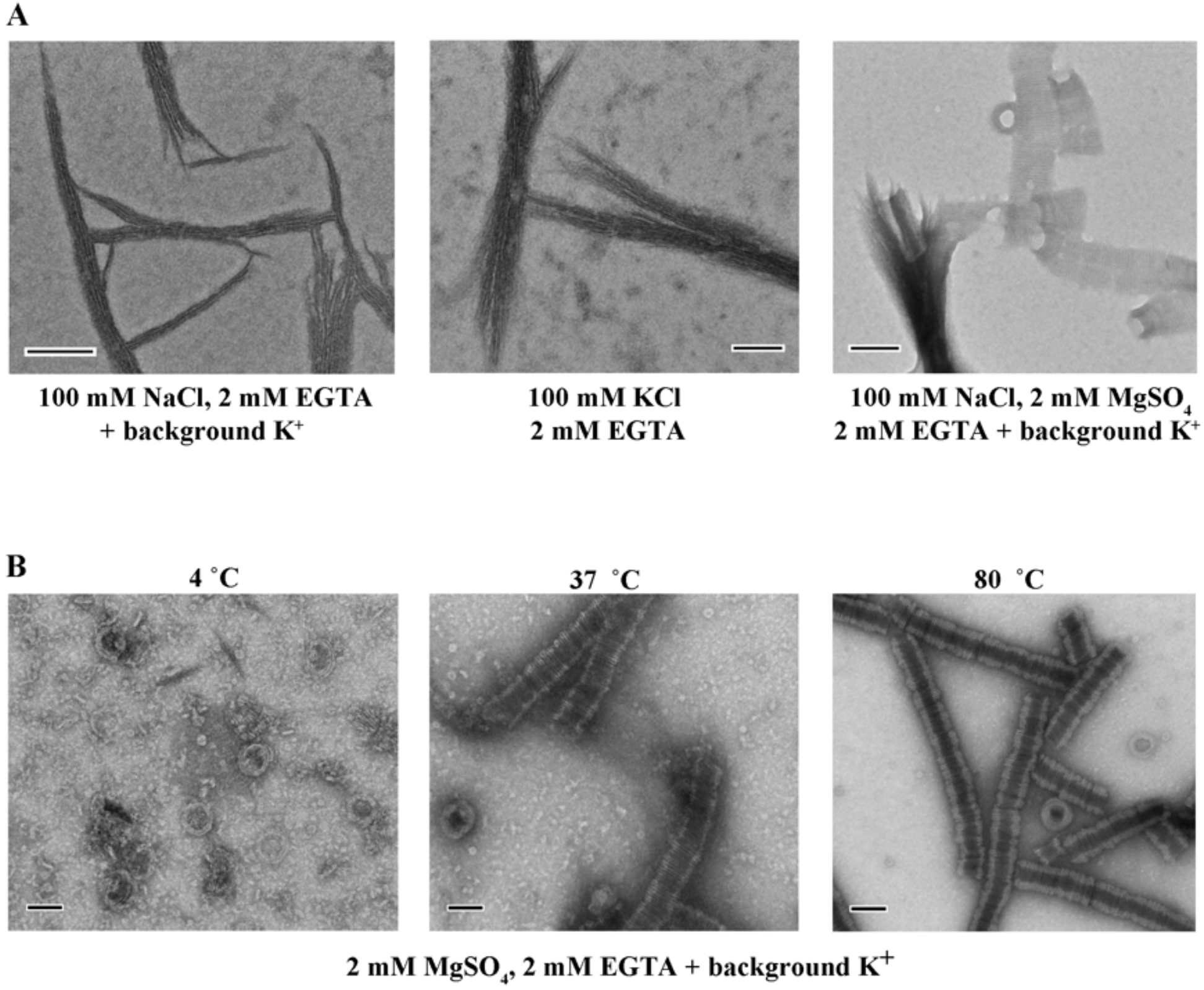
Screening of OdinTubulin assembly conditions observed by EM of negatively stained samples. A) Two morphologies of filaments were observed. Bundles of straight filaments appeared in solutions of monovalent cations (K^+^ or Na^+^). Mixtures of the two forms were observed in mixtures Mg^2+^ with high concentrations of Na^+^. B) The temperature dependence of tubule assembly. Tubules dominated in the presence of Mg^2+^ and absence of Na^+^ at higher temperatures. Scale bar = 100 nm.

### Temperature dependence of filament formation

OdinTubulin forms tubules with a diameter of ∼100 nm in the presence of K^+^/Mg^2+^ at 37 ° or 80°C, with thicker more regular structures formed at the higher temperature (Fig. 6B). Similar to microtubules (*24*), OdinTubulin tubules were not observed at 4 °C, rather immature rings formed that resemble templates for tubule formation (Fig. 6B). The thermostability of the OdinTubulin tubules is consistent with the temperature of the Yellowstone Lower Culex Basin hot spring (∼70 °C) from where Candidatus Odinarchaeota archaeon LCB_4 MAG was sampled (*25*). Next, light scattering was used to monitor the polymerization. OdinTubulin (8 μM) was observed to increase light scattering in the presence of Mg^2+^-containing polymerization buffer without GTP, however the signal was noisy (fig. S7). Including GTP (0.7 mM) in the Mg^2+^-containing polymerization buffer, led to a smooth increase in light scattering over a period of 30 mins, reaching a steady state, consistent with orderly polymerization (fig. S7). Polymerization was temperature dependent. The rate of polymerization increased from 24 °C to 65 °C (Fig. 7A), the temperature limits of the spectrometer.

**Fig. 7.**
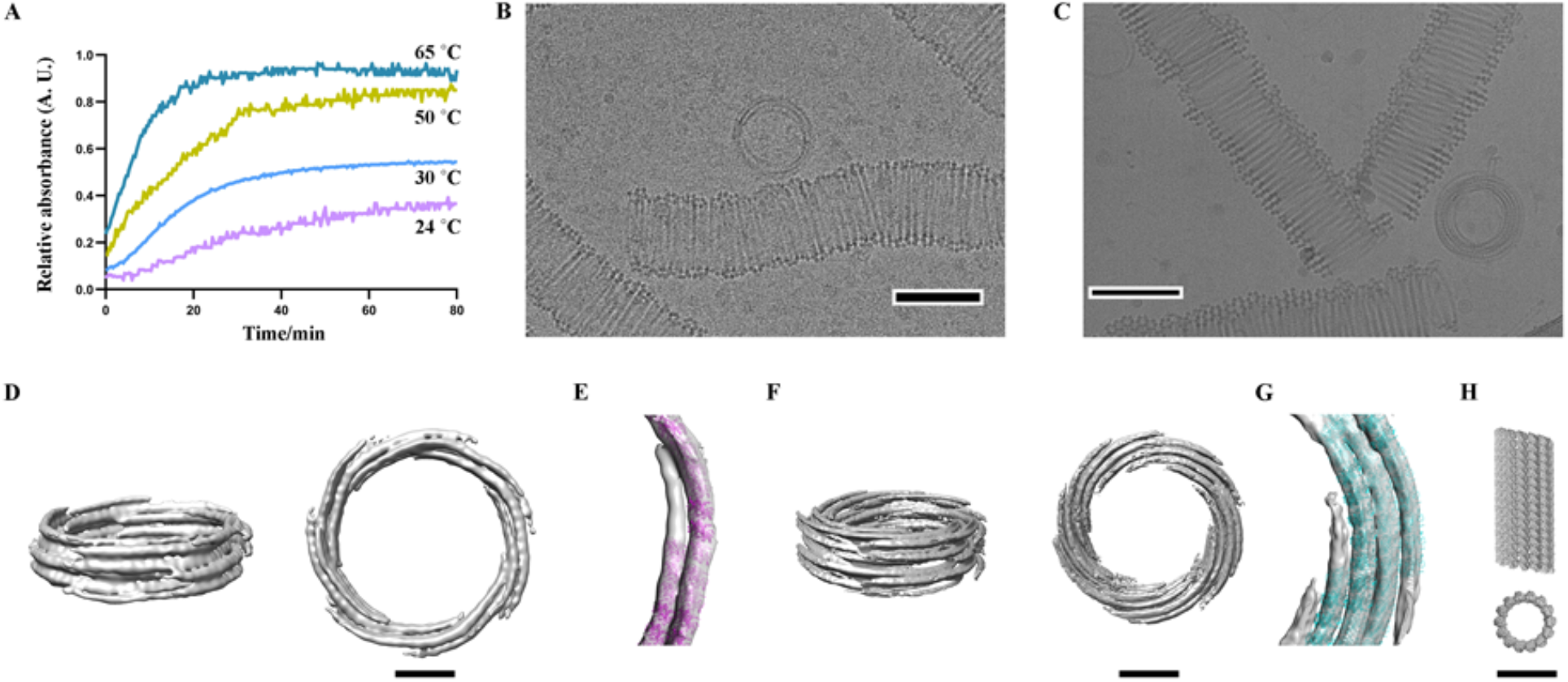
OdinTubulin tubule architecture. (A) OdinTubulin (8 μM) polymerization monitored light scattering at different temperatures. (B) Cryo-electron micrograph of OdinTubulin (40 μM or 60 μM) polymerized at 37 °C and (C), at 80 °C, respectively. Scale bar indicates 100 nm. (D) Two orientations of the 3D reconstruction at 3 nm resolution of OdinTubulin polymerized at 37 °C. (E) The crystal structure fitted into the reconstruction. (F) Two orientations of the 3D reconstruction at 4 nm resolution of OdinTubulin polymerized at 80 °C, and (G) with the fitted model. (H) Two views of the eukaryotic microtubule (*11*). Scale bar in D-H indicates 25 nm.

### OdinTubulin tubule structure

The OdinTubulin tubules are semi uniform but display sufficient homogeneity for calculation of low-resolution reconstructions from cryo-EM images (Fig. 7, B to G). The tubules are constructed from 2-5 layers of short discontinuous curved protofilaments that spiral around the wall of the tubule, approximately perpendicular to the tubule length into which the OdinTubulin crystal structure was placed (Fig. 7, E and G). By contrast, straight protofilaments run along the length of eukaryotic microtubules (Fig. 7H). Tubules assembled at higher temperature (80 °C) appeared more uniform and contained 4-5 layers (Fig. 7, F and G) relative to tubules assembled at lower temperature (37 °C), which typically contained 2-3 layers (Fig. 7, D and E).

We propose that GTP hydrolysis and phosphate release in OdinTubulin protofilaments leads to curving, enabling assembly into tubules, and that the temperature dependence of polymerization can be understood at two levels. At the monomer level, a two-state thermodynamic equilibrium exists, the enthalpic-favoured apo/GDP-bound and the entropically-favored GTP-bound conformations. Elevated temperatures bias the monomer conformation towards the GTP-bound state and protofilament assembly. At the protofilament level, GTP hydrolysis and phosphate release rates will likely increase with temperature, favoring protofilament curving and tubule formation. This second mechanism explains the temperature dependence of the tubule geometries. The lower temperature structures (Fig. 7D), above the assembly temperature threshold, may result from incomplete hydrolysis in the protofilaments during tubule assembly. Odinarchaeota have yet to be isolated, thus the role of the OdinTubulin tubules is unknown. Due to their relatively large diameters (∼100 nm), we speculate that these tubules may use the straight-to-curved protofilament transition to shape membranes. The ∼100 nm diameter of OdinTubulin tubules compares to ∼500 nm diameter of *Candidatus* Prometheoarchaeum syntrophicum MK-D1, a Lokiarchaeon, which is the only Asgard archaeon to be isolated to date (*15, 26*). We hypothesize that duplication of an ancient FtsZ/CetZ gene allowed the OdinTubulin protomer and protofilament to evolve and adopt functions outside cell division.

## DISCUSSION

OdinTubulin forms protomers and protofilaments most similar to eukaryotic microtubules, yet assembles into ring systems more similar to FtsZ (*3*), indicating that OdinTubulin may represent an evolution intermediate between FtsZ and microtubule-forming tubulins. We speculate that enlargement of cell size during eukaryogenesis, may have necessitated the emergence of stiffer tubules to navigate the increasing cellular distances, providing evolutionary pressure that would favor a switch from a malleable tubulin coil geometry to the stiffer parallel protofilament arrangement, seen in microtubules. Such, switches in filament suprastructure architecture, using similar protofilament assemblies, have occurred several times during actin-like and tubulin-like filament evolution (*6, 27*). Gene duplication of the prototypical tubulin gene will have allowed the divergence of α- and β-tubulins to change tubule dynamics, and the straight-to-curved protofilament conformational change repurposed for catastrophe disassembly. Loss of GTP hydrolysis at the N-site in alternate subunits, due to the Glu-to-Lys substitution in the β-subunit (Fig. 4E), may have resulted in relatively less strain, cooperativity and sensing between subunits in the protofilament following hydrolysis, extending the transient stability of the straight form of the protofilaments in the early microtubule. Another gene duplication event allowed the emergence of γ-tubulin as a nucleation complex in a parallel scenario to actin gene duplication in the emergence of the ARP2/3 actin-filament nucleating complex (*28*).

Further evidence for OdinTubulin representing a record of the prototypical tubulin prior to the evolution into microtubule-forming tubulins can be found in sequence analysis. Comparison of a hybrid human α/β-tubulin sequence, which includes the interface residues at the E-site from both α-tubulin and β-tubulin, increased the identity with OdinTubulin from 35% for α-tubulin and β- tubulin to 38% for the hybrid sequence, indicating that OdinTubulin represents a reasonable model of eukaryotic tubulin prior to the gene duplications. Thus, tubulin is an example in evolution, in which gene duplication coupled with sequence variation, without significant structural change to the core protein component (the tubulin protomer), gave rise to a novel complex protein machine, the microtubule. The microtubule is essential to eukaryotic chromosome segregation and its emergence was likely a key event in eukaryogenesis. In summary, OdinTubulin appears to have the characteristics of a primordial tubulin before the transition into the eukaryotic microtubule-forming tubulins.

## MATERIALS AND METHODS

### Protein expression and purification

The *Escherichia coli* codon optimized gene encoding the Odinarchaeota tubulin protein (OLS18786.1) was synthesized and placed in the pSY5 vector which encodes an N-terminal HRV 3C protease cleavage site and 8-histidine tag (*14*). The OdinTubulin mutation (H393D) was introduced using the Q5 Site Directed Mutagenesis Kit (New England BioLabs) according to the manufacturer’s protocol. Plasmids were transformed into *E. coli* (DE3), the cells grown to a density of OD_600_ = 0.8 and the protein expressed by induction with 0.5 mM isopropyl-D-1-thiogalactopyranodside (IPTG) at 18 °C overnight. After centrifugation, cell pellets were resuspended in binding buffer (20 mM HEPES, 500 mM NaCl and 1 mM TCEP, pH 7.5), supplemented with Triton X-100 (0.01%), protease inhibitor cocktail (Set III, EDTA-free, Calbiochem) and benzonase (2 μl of 10,000 U/μl, Merck) or in EM binding buffer (100 mM PIPES, 500 mM NaCl, 50 mM imidazole, 10 mM MgSO_4_, 2 mM EGTA, pH 6.9). Cells were lysed using an ultrasonic cell disrupter Vibra-Cell (Sonics). The protein was purified from the clarified supernatant using a Ni-NTA affinity chromatography column (HisTrap FF GE Healthcare) with binding buffer and eluted through on column cleavage with HRV 3C protease. Affinity purified protein was further purified by size-exclusion chromatography (16/60 Superdex 75 PG, GE Healthcare) in the gel filtration buffer (20 mM HEPES, pH 7.5, 150 mM NaCl, 1 mM TCEP, or for EM samples: 100 mM PIPES, pH 6.9, 150 mM NaCl, 1 mM MgSO_4_, 2 mM EGTA and 50 μM GTP). Pure protein containing fractions were identified by SDS–PAGE, pooled and concentrated with 2000-10000 MWCO Vivaspin concentrators (Vivascience), and flash frozen in liquid nitrogen in small aliquots, or used freshly.

### Crystallization, structure determination, model building and refinement

Native OdinTubulin and H393D crystallization trials, at 5-15 mg/ml in 20 mM HEPES, pH 7.5, 150 mM NaCl, 1 mM TCEP) were performed using the sitting-drop or vapour-diffusion methods with a precipitant solution (1:1) at 293 K. Native OdinTubulin crystals were formed in 0.1 M Bis-Tris, pH 7.5, 25 % w/v PEG 3350. These crystals diffracted X-rays poorly to 4 Å, however the resulting data set was amenable to successful molecular replacement using the *Sus scrofa* β-tubulin structure as a search model (PDB 6o2r, chain K) (*18*). Mutational analysis, of crystal contacts, identified a single amino acid substitution (H393D) that had improved diffraction to 2.5 Å, but did not alter protofilament packing, however the nucleotide-binding site showed partial occupancy (7EVH, fig. S8 and table S1). Soaking of these crystals with GTP increased the diffraction limit and sharpened the electron density suitable for unambiguous structure determination (table S1). Crystals were frozen in the mother liquor. X-ray data were collected on RAYONIX MX-300 HS CCD detector on beamline TPS 05A (NSRRC, Taiwan, ROC) at λ = 1.0 Å or on BL41XU (λ = 1.0 Å) of SPring-8 on a Pilatus 6M detector.

Data were indexed, scaled, and merged following standard protocols (*16*). Molecular replacement and refinement was carried out using PDB 6o2r chain K as the search model using standard methods to solve the 1.62 Å structure 7EVB (table S1) (*16*). The identity of the bound nucleotide was assessed by refinement of GTP, GDP or a combination of GTP and GDP in the nucleotide-binding site. Subsequent soaks and alternate crystal structures are detailed in table S1.

### Polymerization assay

Polymerization of native OdinTubulin or H393D (8 μM) was induced by the addition of GTP (0.7 mM) in K-PIPES buffer (100 mM PIPES, pH 6.9, 0.5 mM EGTA, 0.5 mM MgSO_4_, 10% (v/v) glycerol), total volume of 100 μl at various temperatures. Absorption at 340 nm was used to measure an increase in light scattering consistent with polymerization in 96-well, clear, flat-bottomed plates (Corning, Nunc). The plates were equilibrated (30 min) to the appropriate temperatures prior to the assays. Changes in absorbance were monitored with an Infinite M Nano^+^ plate reader (Tecan).

### Interference Reflection Microscopy (IRM)

Glass cover slips and slides were cleaned 30 min in Hellmanex III (2% in water) at 60 °C with sonication, and rinsing in ultrapure water. Cover slips were dried using nitrogen gas flow. *In vitro* polymerization assays were performed using flow chambers with dimensions of 3 × 20 × 0.07 mm (width × length × height) that were assembled with double-sided tape as the spacer from 20 × 20 mm cover slip and slide. Brain tubulin elongation assay: Seeds (in the mix) were elongated with a mix containing 15 μM of tubulin at 30 °C in IRM buffer: 100 mM Pipes-K pH 6.9, 0.5 mM MgSO_4_, 0.5mM EGTA supplemented with 0.7 mM GTP, an oxygen scavenger cocktail (20 mM DTT, 3 mg/ml glucose, 20 μg/ml catalase and 100 μg/ml glucose oxidase) and 0.25% methyl cellulose (1,500 cP, Sigma). OdinTubulin was similarly polymerized by addition of 0.7 mM GTP in the IRM buffer at the protein concentrations and cation conditions indicated in Fig. 3.

Non-labelled microtubules and non-labelled OdinTubulin filaments were imaged with IRM on an epifluorescence microscope (Eclipse Ti2, Nikon). The samples were illuminated with a SOLA Light Engine (Lumencor) through a cube equipped with a monochromatic filter at 520 nm, a 50/50 dichroic mirror and a × 60 numerical aperture 1.49 TIRF objective. The microscope stage was kept at 30 °C using a warm stage controller (LCI). Images and movies were captured using an Orca flash 4LT camera (Hamamatsu) every 5 s for 30 min.

### Sequence and structure analyses

Tubulin, FtsZ and CetZ structures were aligned in using PDB codes 1rlu, 1w5e, 2vam, 2vaw, 3zid, 4b45, 4e6e, 4ffb, 5jco, 5mjs, 5n5n, 5ubq, 5w3j, 6e88, 6rvq, and 6unx. The resulting alignment was subjected to phylogenetic analysis using published methods (*16*). Structure comparisons were carried out using the Dali server (http://ekhidna2.biocenter.helsinki.fi/dali/).

### MD simulations

The all-atom molecular dynamics simulations were performed by the software GROMACS 2019 (*29*). The Amber ff14SB force field (*30*) and TIP3P (*31*) water model were used for the protein molecule and solvent. The force field parameters for the GTP were taken from (*32*). The atomic coordinates of the tubulin subunits were taken from the Protein Data Bank (PDB code 6o2r) (*18*). The protein atoms were solvated in the truncated octahedral water boxes (with 28413 water molecules). Na^+^, K^+^, and Cl^-^ were added to neutralize the systems and to simulate the salt concentrations of [NaCl]=10 mM and [KCl]=150 mM. Covalent bonds involving hydrogen atoms were restrained by the LINCS algorithm (*33*). The system was firstly minimized by 50000 steps using the steepest descent method, and then equilibrated for 0.1 ns in the canonical ensemble (NVT) and another 1.0 ns in the isothermal-isobaric ensemble (NPT). After the equilibration simulations, the production simulations with the length of 1.0 μs were conducted. The temperature and pressure were controlled at 298.0 K and 1.0 atm, respectively. We performed seven independent simulations with different initial atom velocities. In calculating the occupancies, snapshots of the first 200 ns in each of the MD trajectories were omitted. MDTraj was used for analysis (*34*).

### Negative Staining of Odin Tubulin

Negatively stained GTP-induced tubules were observed using a Hitachi-H7600 transmission electron microscope (Institute for Advanced Research, Nagoya University). Thawed OdinTubulin protein was diluted to final concentration of 40-60 μM in pre-warmed polymerization K-PIPES buffer as previously described for eukaryotic tubulin (*35*) and polymerized for 10-20 min using 2 mM GTP (Sigma-Aldrich). 2.5 μl of the mixture was applied onto glow discharged grid STEM100Cu elastic carbon grids (Ohkenshoji co., Ltd) and absorbed for 1 min. The sample was blotted with filter paper and then 2% uranyl acetate solution (5 μl) was applied. After 1 min, the grids were blotted again and allowed to dry overnight.

### Cryo-EM grid preparation

Molybdenum R1.2/1.3 and R2/2 200-mesh grids with a holey carbon support film (Quantifoil, Jena, DE) were glow discharged for 40 s under high vacuum shortly before sample application. OdinTubulin protein (40-60 μM) was polymerized for 10-20 minutes at 37 °C and 80 °C using an adapted protocol for eukaryotic tubulin (*35*) by adding temperature equilibrated K-PIPES buffer and 2 mM GTP (Sigma-Aldrich). 2.5 μL of the polymerized sample was applied to the glow-discharged grids, blotted for 0.5–1.5 s and plunge-frozen with an EM GP plunge freezer (Leica Microsystems) operated at room temperature at 90% humidity.

### Cryo-EM data acquisition and image processing

Frozen molybdenum cryo-EM grids were initially screened on a JEOL JEM-3000SFF electron microscope (Cellular and Structural Physiology Institute, Nagoya University) operated at 200 kV at minimal dose system. Images were recorded using a K2 Summit camera with exposure settings of 1.35 Å/pixel size. Grids were subsequently imaged on a Titan Krios (FEI), at the Institute for Protein Research, Osaka University, equipped with the FEG operated at 300 kV and a minimal dose system. Imaging was performed using the EPU software (FEI) or SerialEM software (Nexperion) attached to the Titan Krios. Images of OdinTubulin incubated at 37 °C were recorded at nominal magnification of 47,000, without using objective aperture, nominal defocus range of - 2.0 to -2.6 μm with a dose rate of 40.05 e^-^/Å^2^ and exposure time of 2.42 s. Images were recorded using a Falcon III detector (FEI) at a pixel size of 1.45 Å/pixel and a frame rate of 60 frames/individual images. For incubation at 80 °C, two sessions of data collection at a magnification of 64,000, without objective aperture, defocus range of -2.0 to -2.4 μm with a dose rate of 50 e^-^/Å ^2^ and exposure time of 5.21 seconds were used. Images were recorded with a K3 summit detector (FEI) in counting mode at a pixel size of 1.11 Å/pixel and a frame rate of 58 frames per image.

1217 (for 37 °C) and 7600 (for 80°C) raw movies were collected and processed in RELION 3.0/3.1.1 (*36*). Drift was motion corrected with MotionCor2 (*37*) and the CTF for each micrograph was estimated with CTFFind-4.1 (*38*). Micrographs with good observed CTF estimations were selected for further processing. Tubules were picked manually using EMAN2 e2helixboxer (*39*) and extracted in RELION 3.0/3.1.1 with a 4 × 4 binning (box size of 250 × 250 pixels). Particles from 2D classes displaying clear and similar structure and radius were selected. The helical pitch of the coil was determined using the 2D classes. 3D classification was performed with a wide range of initial helical parameters using the helical pitch of the coil as a restriction. The 3D classes which had consistent projections with 2D classes and top views of the filaments were selected. Initial 3D reference models were prepared using RELION toolbox kit cylinder. Two rounds of 3D classification were performed. 3D refinement was performed with a reference model low pass filtered at 40 Å with solvent mask. Particles were re-extracted with a 2 × 2 binning (box size of 500 × 500 pixels) and the final 3D refinement was performed.

### Model fitting

In the crystals, the protofilaments are unable to bend due to the crystal packing. However, superimposing the apo-OdinTubulin (7EVG) onto two adjacent subunits from the GTP-bound protofilament (7EVB), via the C-terminal domains, led to a curved model that could be fitted into the EM density.

## Acknowledgements

We thank the Synchrotron Radiation Protein Crystallography Facility of the National Core Facility Program for Biotechnology, Ministry of Science and Technology and the National Synchrotron Radiation Research Center, a national user facility supported by the Ministry of Science and Technology, Taiwan, ROC, and the SPring-8 Synchrotron, Japan. EM screening and data collection was supported by the Japan Agency for Medical Research and Development (AMED) Grant Number JP20am0101074 (A.O.) and the Collaborative Research Program of Institute for Protein Research, Osaka University (CENCR-20-20). We thank Esra Balıkçı for technical support.

## Author contributions

C.A., S.A., A.N., L.B. and R.C.R. conceived experiments and analyzed data. C.A., S.A., and L.T.T performed biochemical experiments. C.A., L.T.T and R.C.R. conducted X-ray experiments. J.G. performed I.R.M. experiments and W.L. conducted MD simulations. S.A., A.N., A.O. and K.H. conducted EM experiments. A.N., K.F., L.B. and R.C.R. supervised the work. R.C.R. wrote the manuscript. All authors edited the manuscript.

## Data and materials availability

The atomic coordinates and structure factors have been deposited in the Protein Data Bank under the accession codes: 7EVB-D, 7EVG-L and 7F1A-B. All other data are available in the main text or the supplementary materials.

## Supplementary Materials

**Fig. S1.**
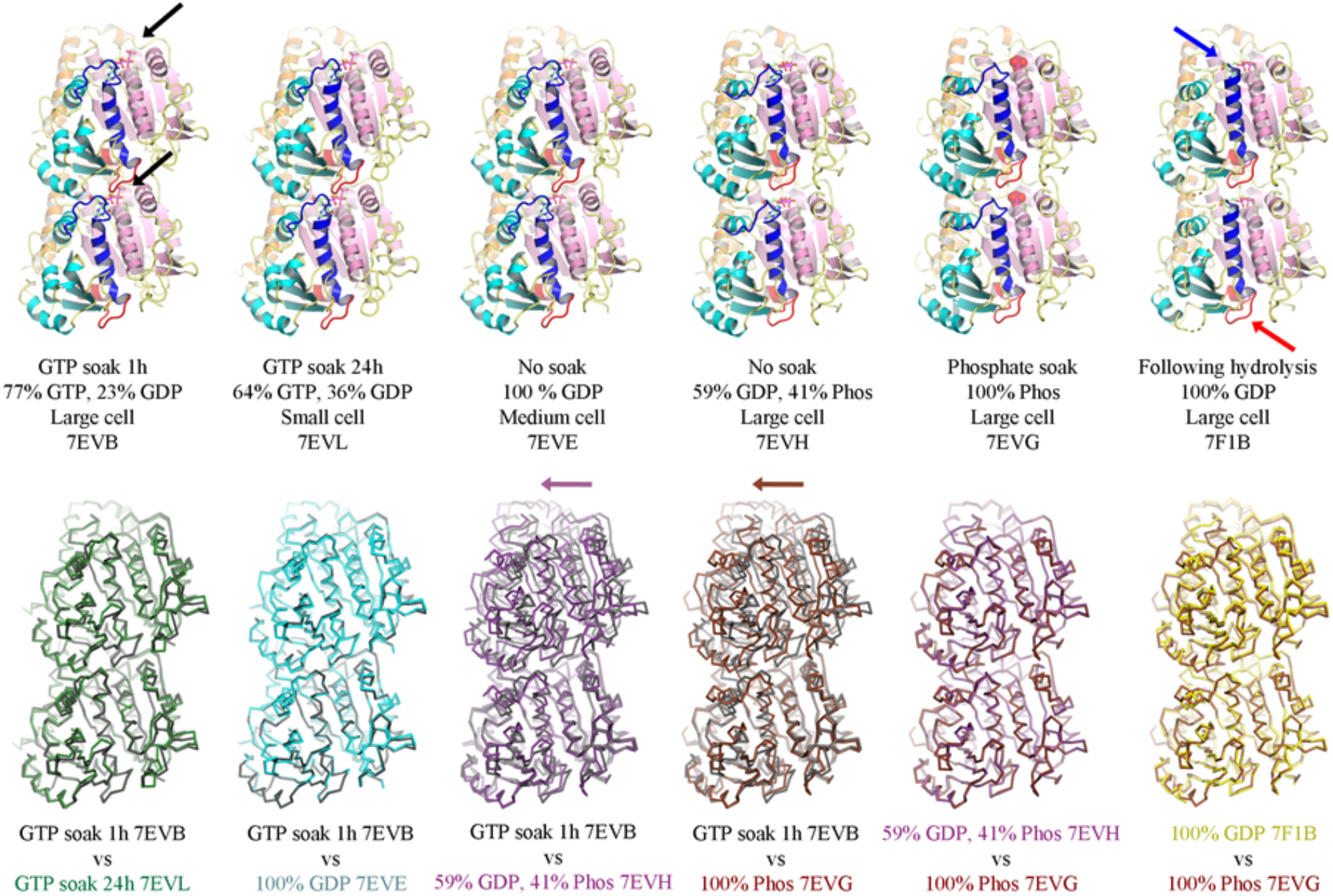
Six classes of OdinTubulin crystals and structures from this study. Top row: Two copies of the structures from adjacent asymmetric units in the crystals, which form protofilaments (7EVB, 7EVL and 7EVE) or pseudo-protofilaments (7EVH, 7EVG and 7F1B). Secondary structure elements are colored by domain: N-terminal (pink), intermediate (cyan), and C-terminal (orange). The nucleotide sensor motif (red and blue, indicated by arrows on the right structure) lies within the intermediate domain. The nucleotides are shown as sticks and highlighted by black arrows in the left structure. Free phosphate is shown as spheres. Lower row: superimposition of structures indicating two arrangements. The protofilament forming structures (7EVB, 7EVL and 7EVE) and the pseudo-protofilaments forming structures (7EVH, 7EVG and 7F1B). Arrows indicate the shift between the protofilament- and pseudo-protofilaments-forming structures. The shift away from the protofilament arrangement likely occurs due the curved form of the protofilament being incompatible with the translational symmetry of the crystal, leading to a translation, rather than a curving in the restrained crystal environment. The other five structures from soaked crystals (7EVC, 7EVD, 7EVI, 7EVK and 7F1A) adopt the large cell protofilament crystal form, similar to 7EVB.

**Fig. S2.**
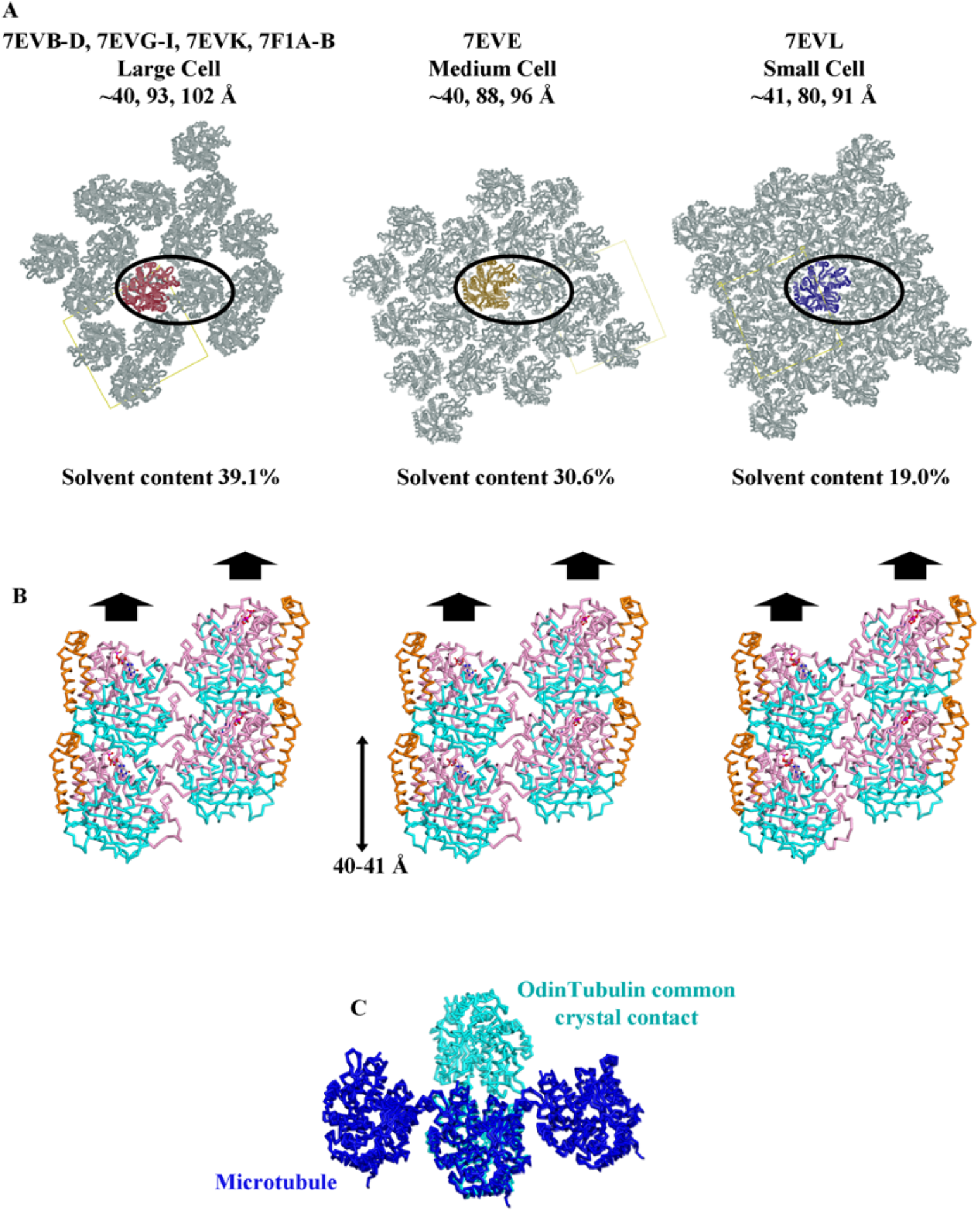
Packing in OdinTubulin crystals. (A, B) The three classes of cell dimension, all in space group P2_1_2_1_2_1_. In each arrangement, the crystal forms from the staggered association of two parallel protofilaments (circled in A), or pseudo protofilaments, with different solvent contents. (A) top view, (B) side view with the bold arrows indicating the protofilaments or pseudo protofilaments. In B) the domains are colored N-terminal (pink), intermediate (cyan), and C-terminal (orange). (C) Comparison of the staggered association of two parallel OdinTubulin protofilaments (cyan) from the crystals with three adjacent subunits around a microtubule (blue). We do not interpret the staggered association of the two parallel OdinTubulin protofilaments in the crystal packing to have physiological relevance, since the arrangement it is not found in the EM structures.

**Fig. S3.**
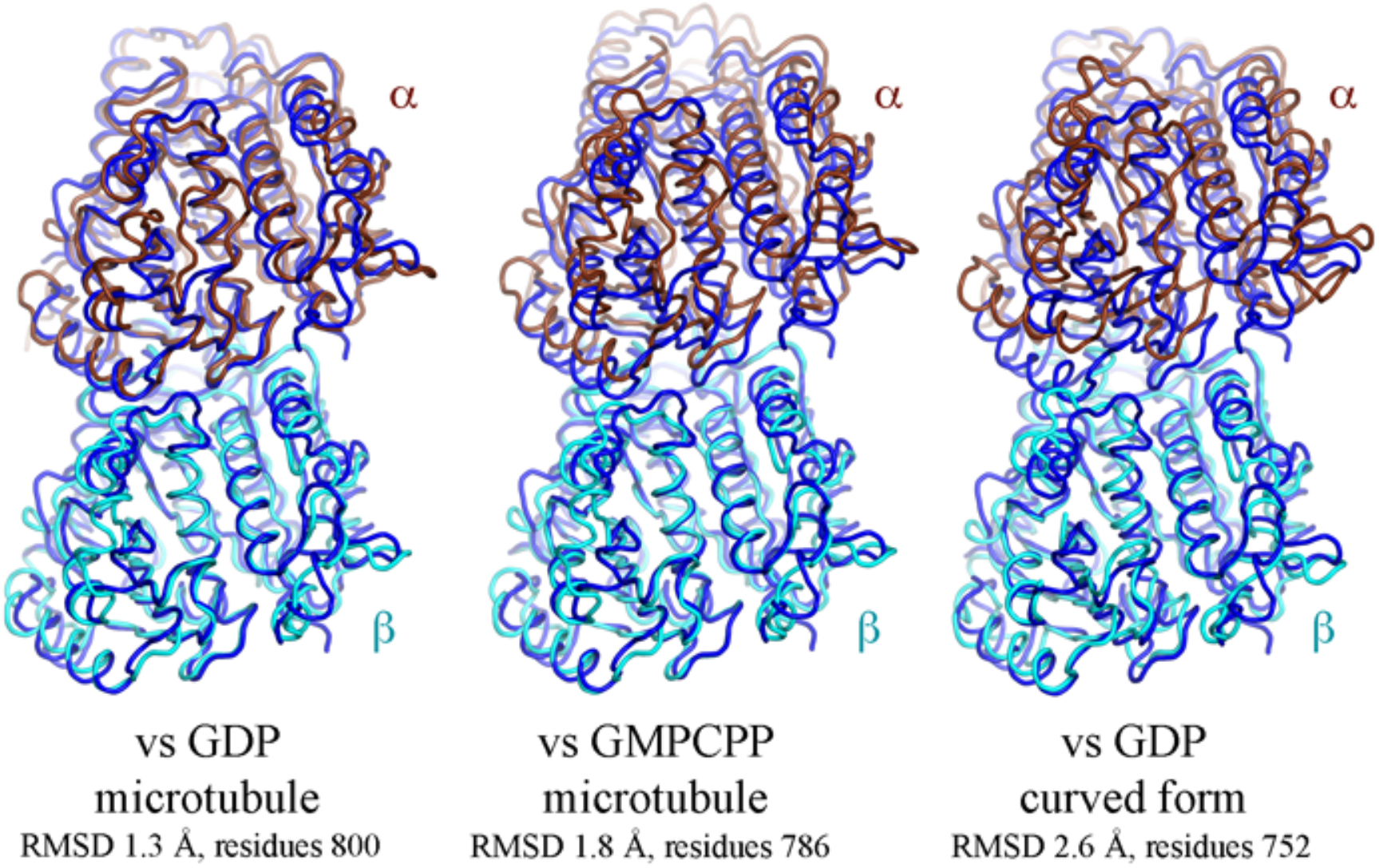
Comparison of OdinTubulin protofilaments with eukaryotic tubulin. Superimposition of the two GTP-bound OdinTubulin symmetry-related subunits from the crystal packing (dark blue) onto two subunits of eukaryotic tubulin from the GDP-bound microtubule (PDB 6o2r) (Eshun-Wilson et al., 2019), the guanosine-5’-[(αβ)-methyleno]triphosphate (GMPPCP)-bound microtubule (PDB 6dpu) (Zhang et al., 2018), and the stathmin-bound curved protofilament (PDB 4iij) (Prota et al., 2013). α- and β-tubulins are shown in brown and cyan, respectively. The RMSD statistics indicate the structural similarity of the α/β-tubulin heterodimer to the pair OdinTubulin subunits. See table S3 and movie S1 to S3.

**Fig. S4.**
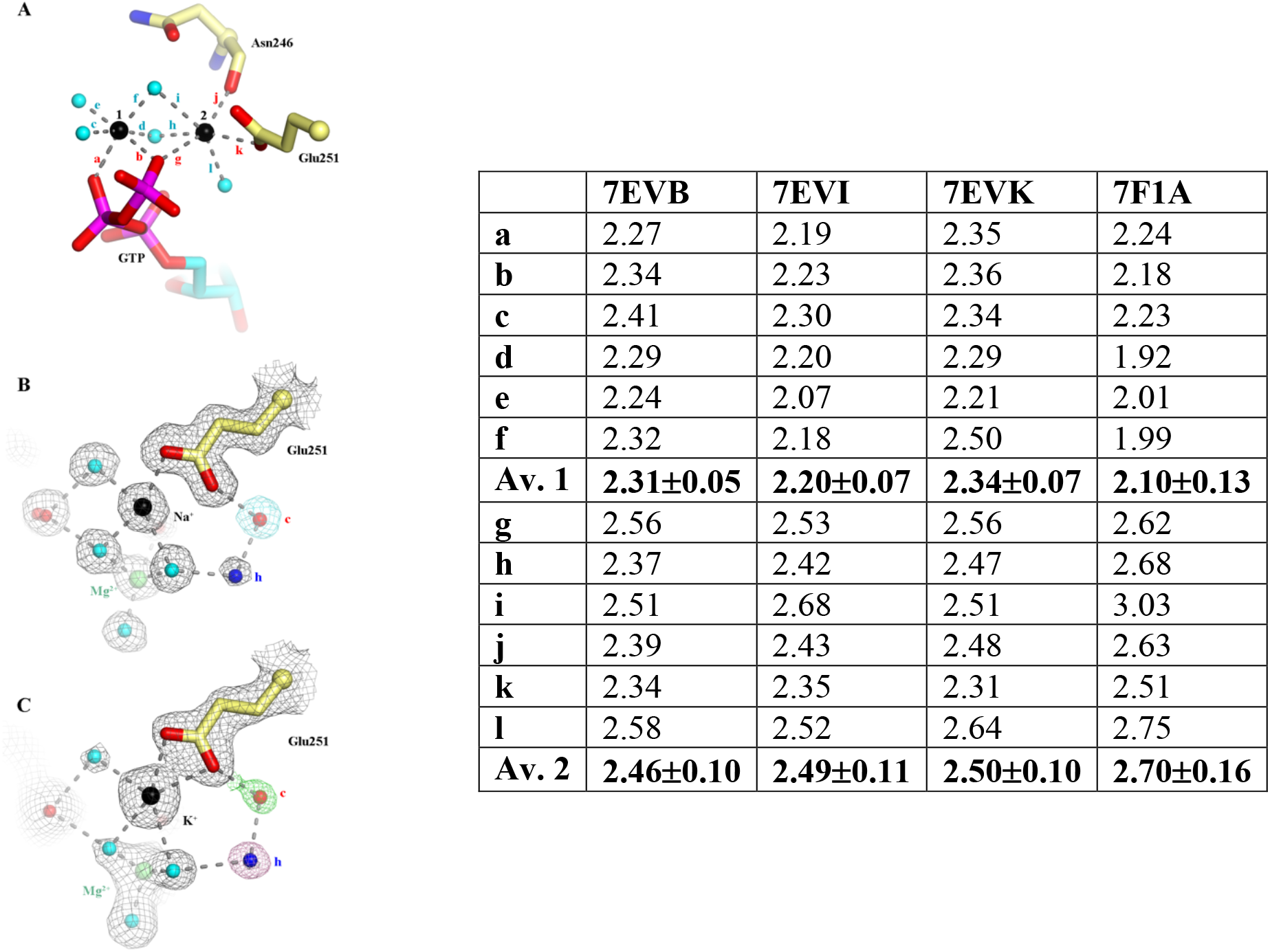
The identity of the two cations. The octahedral geometry (A) and bond lengths (table) of the cations. In structure 7EVB, in 200 mM sodium acetate, both cations are likely to be sodium, or have mixed occupancy resulting from background cations. The average bond lengths (Av.) of cation 1 are significantly shorter than for ion 2. Structure 7EVI was soaked with 2 mM MgCl_2_ and 200 mM sodium acetate (1 h), which reduced the bond lengths for cation 1 to 2.2 Å, typical for Mg^2+^, while those for cation 2 remained unaffected. Structure 7EVK was soaked with 2 mM EGTA, 100 mM KCl, 200 mM NaCl (1 h) to remove any divalent cations and to determine whether the bond lengths increased due occupancy by K^+^. The bond lengths slightly increased, indicating that both cation-binding sites can accept monovalent cations. Structure 7F1A was soaked with 2 mM MgCl_2_ and 200 mM KCl (1 h) and had bond lengths consistent with Mg^2+^ and K^+^. Thus, the Mg^2+^ is the preferred cation at site 1, and site 2 is a monovalent cation binding site, preferentially occupied by K^+^/Na^+^, under the conditions tested. (B) The relationship between Na^+^ in site 2 (7EVI) and the proposed hydrolytic and hydrogen-receiving waters, c and h. The atoms are surrounded by the OMIT map contoured at 1 σ (grey) and highlighted for atom c (cyan). (C) Similar representation for the Mg^2+^/K^+^ soak (7F1A). The K^+^ coordination appears to be pentagonal bipyramidal, the coordination increased by E251 providing double coordination. This has ramifications for the occupancies of the c and h waters. h is barely visible, and c not visible in the OMIT map (grey), but refine in the 2Fo-Fc map (1 σ), green and pink, respectively. Thus, the exact positioning of E251 effects the c and h waters, consistent with its proposed role in hydrolysis.

**Fig. S5.**
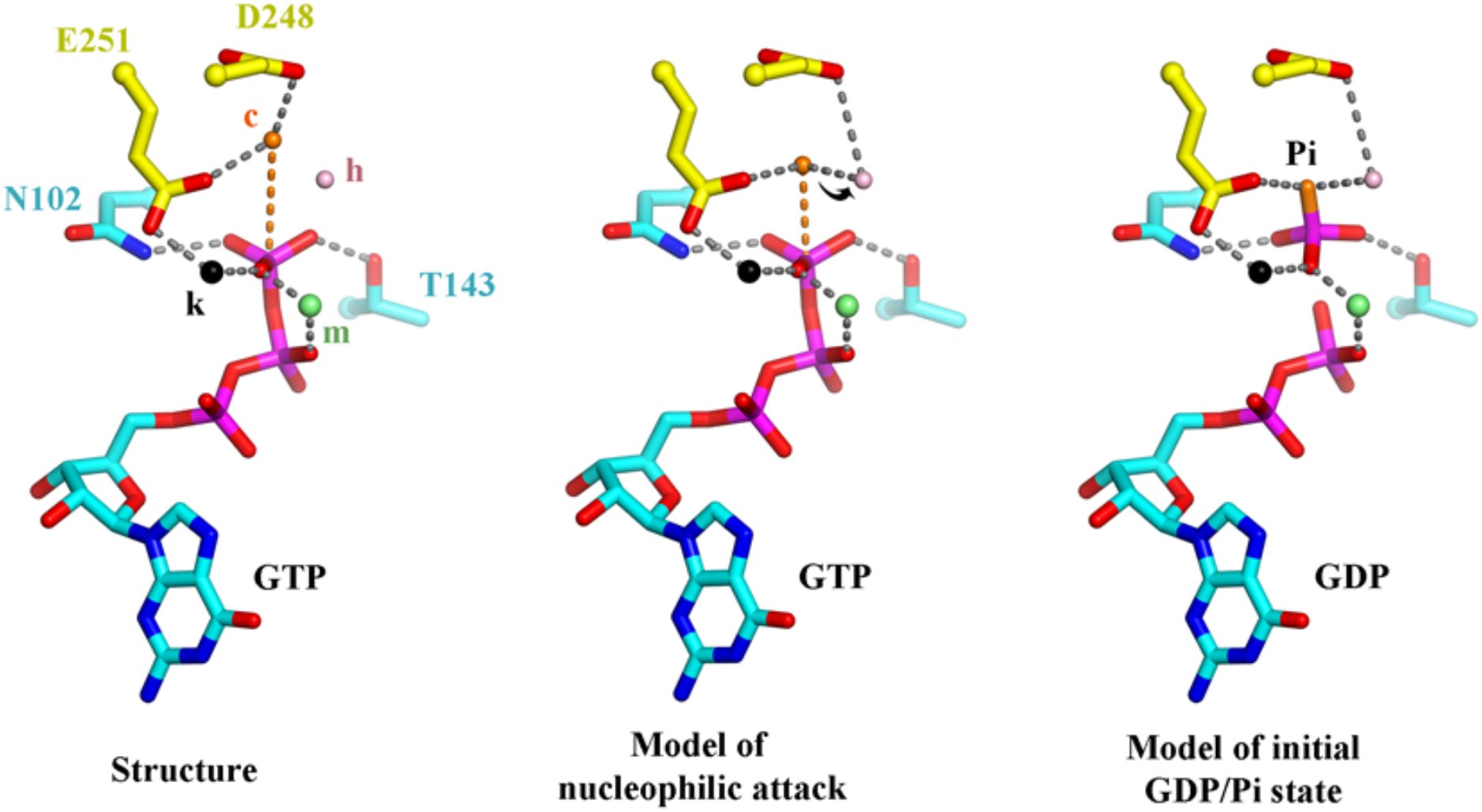
Hypothetical model of GTP hydrolysis in OdinTubulin. Left, Structure 7EVI is shown with the residues and ions that stabilize the GTP γ-phosphate. c, hydrolytic water; h, hydrogen ion receiving water; m, magnesium ion; k, potassium or sodium ion. In the crystal structure water c is 4.1 Å from the GTP γ-phosphate phosphorous atom. Middle, to initiate hydrolysis the hydrolytic water (orange) is required to approach the GTP γ-phosphate phosphorus atom, along the orange dashed line, likely losing a hydrogen ion to the hydrogen ion receiving water (pink), indicated by the arrow. Left, after hydrolysis the dissociated γ-phosphate ion will receive a hydrogen ion, possibly from a surrounding water molecule or from Thr143. The long distance of water c is 4.1 Å from the GTP γ-phosphate phosphorous atom may be due to two reasons. Firstly, a trivial reason that the H393D mutation may slightly increase the distance (fig. S8). Secondly, the extended distance may be part of the cooperative conformational changes within a protofilament. We hypothesize that protofilaments initially as straight protofilaments. Stochastic fluctuations in the water c position may then initiate hydrolysis in one or more subunits, which in turn may be propagated by the allosteric motion of the nucleotide sensor motif to produce concerted hydrolysis and conformational change throughout the protofilament.

**Fig. S6.**
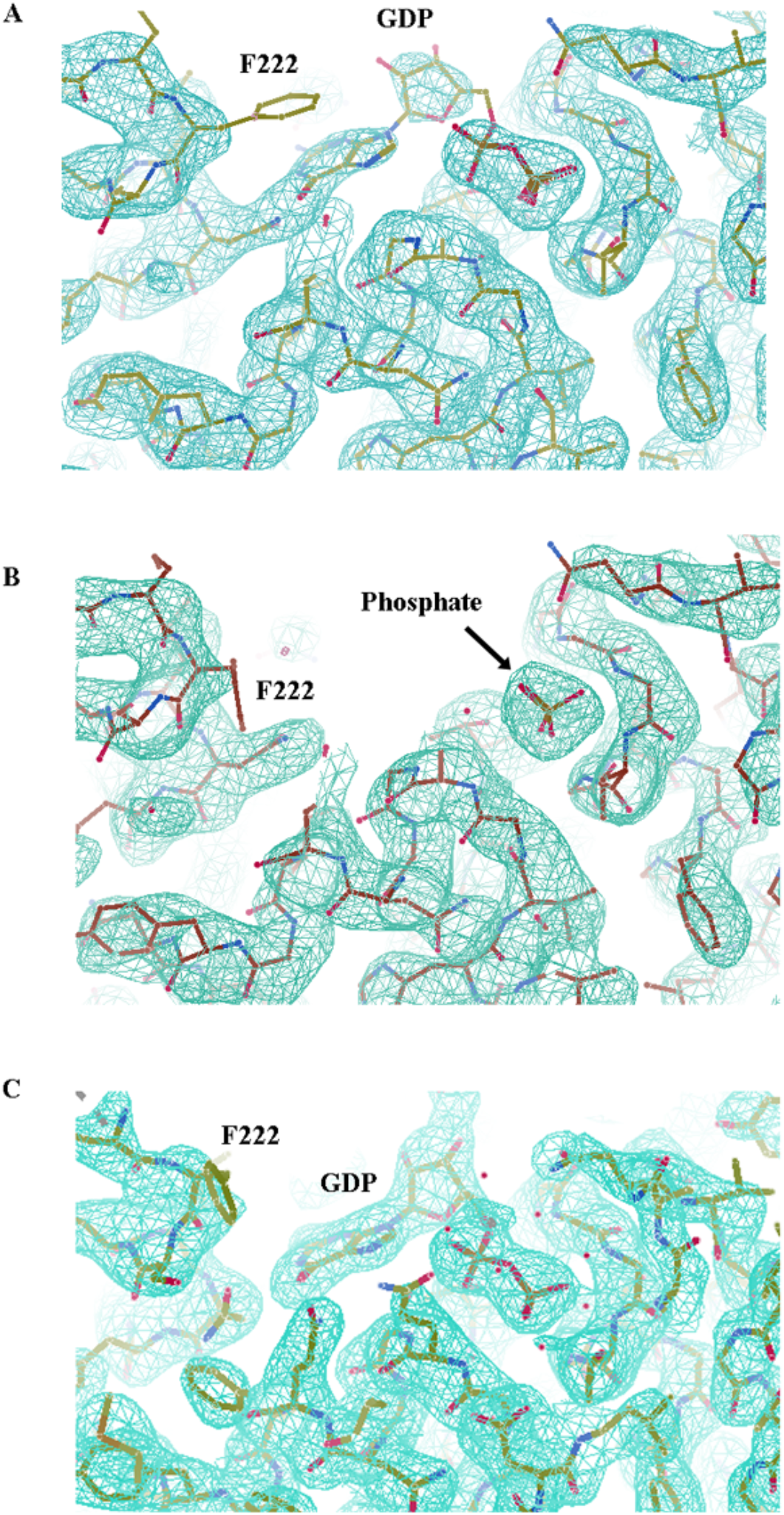
The second conformation of OdinTubulin. The OMIT maps contoured at 1 σ for the alternate conformation of OdinTubulin bound to (A) 60% GDP and 40% phosphate (7EVH), (B) 100% phosphate (7EVG) or (C) 100% GDP (7F1B) from hydrolysis reaction carried out within the crystal. Phe222, which forms phi:phi stacking with nucleotide in the other conformation (Fig. 2B) is disordered in these structures, providing the mechanism for nucleotide dissociation.

**Fig. S7.**
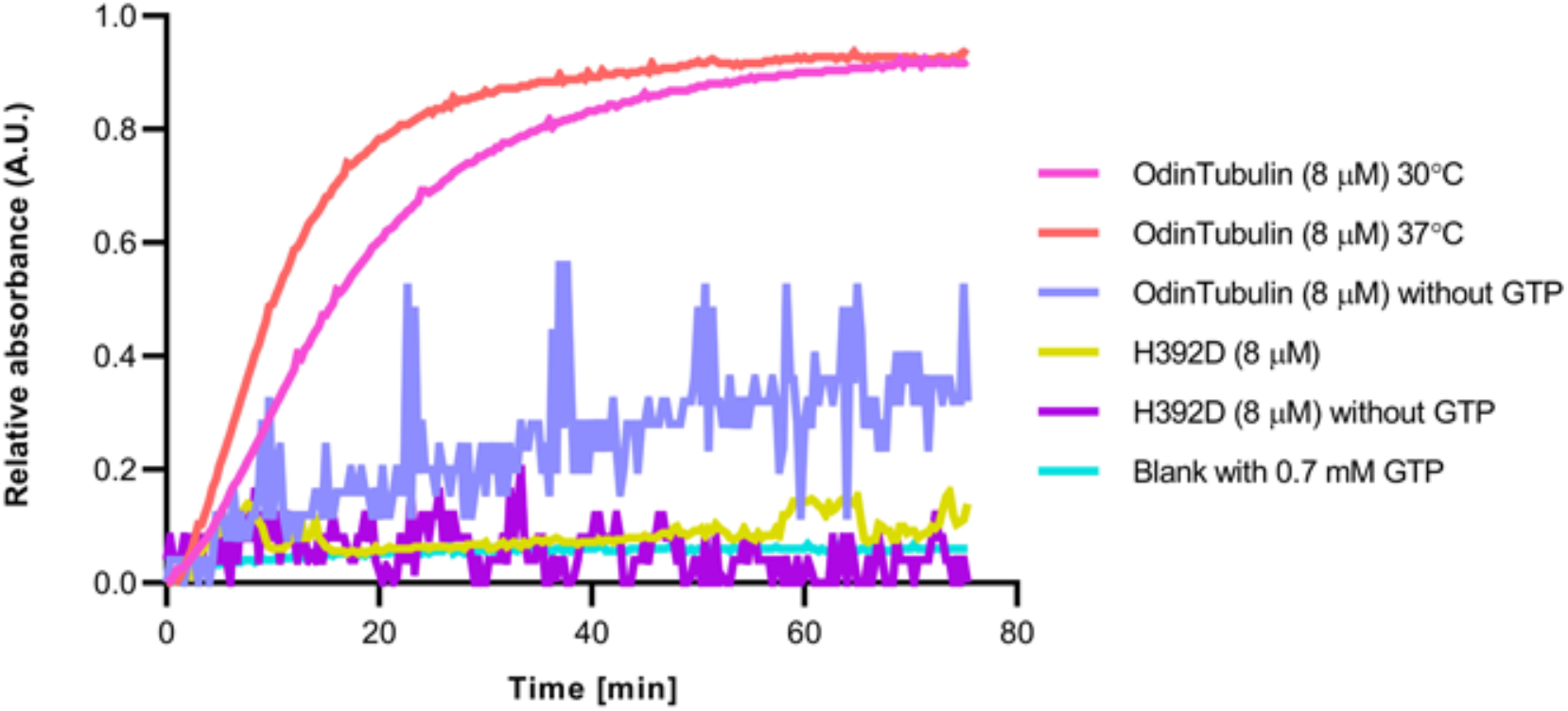
Polymerization of OdinTubulin. Light scattering profiles for native and H393D mutant OdinTubulin (8 μM) on polymerization with and without GTP.

**Fig. S8.**
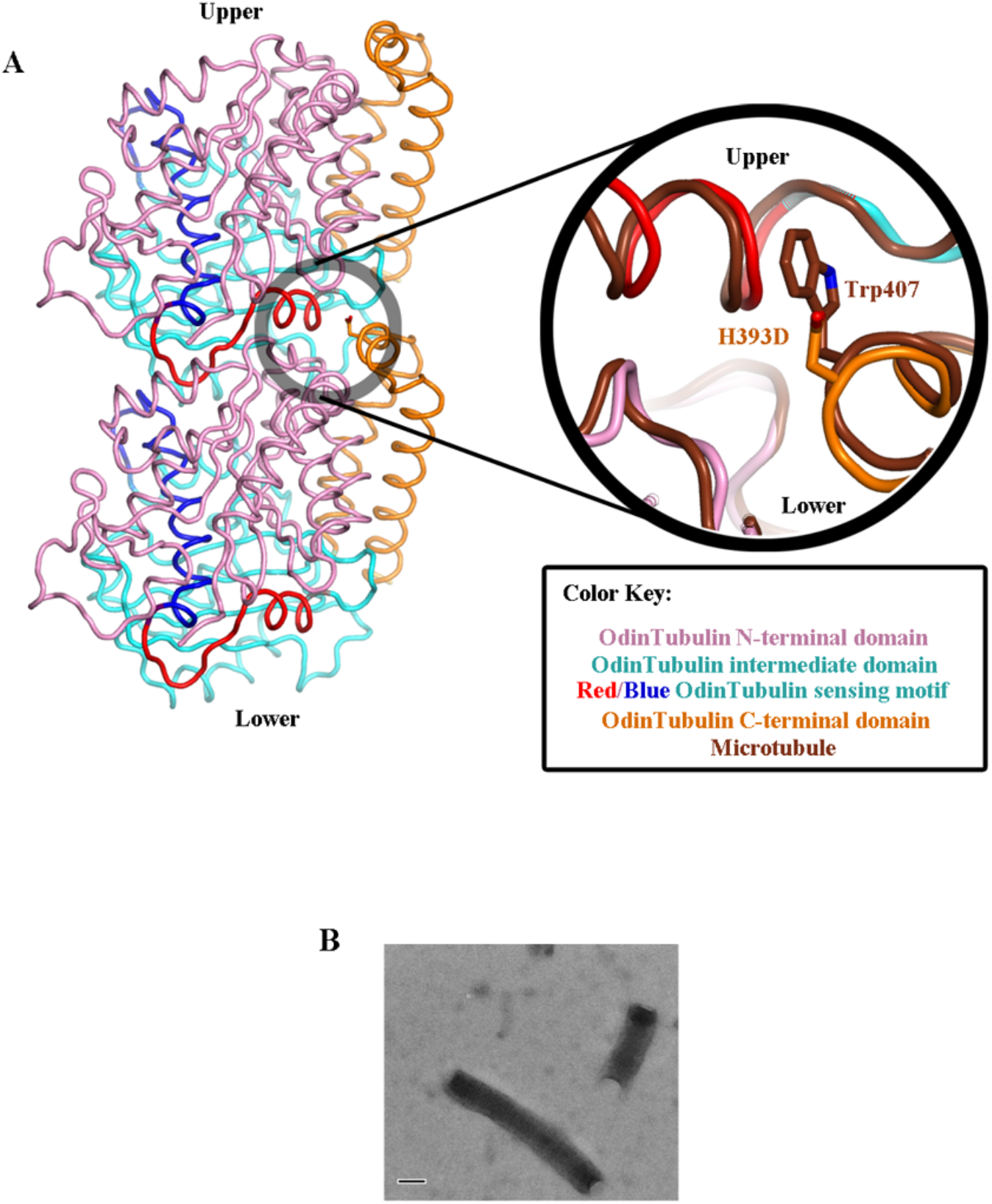
The OdinTubulin H393D mutation. A) The H393D mutation in OdinTubulin lies at the edge of the packing between two subunits within the protofilament. The enlargement demonstrates that this mutation does not significantly affect the packing between the subunits in comparison to the microtubule protofilament packing (brown). B) Light scattering showed reduced assembly of the H393D OdinTubulin mutant (fig. S7). Observation of negatively stained samples H393D OdinTubulin mutant tubules by EM demonstrated that tubules of the mutant form in a similar morphology to the wild type protein, however the assembly efficiency is lower.

**Table S1.**
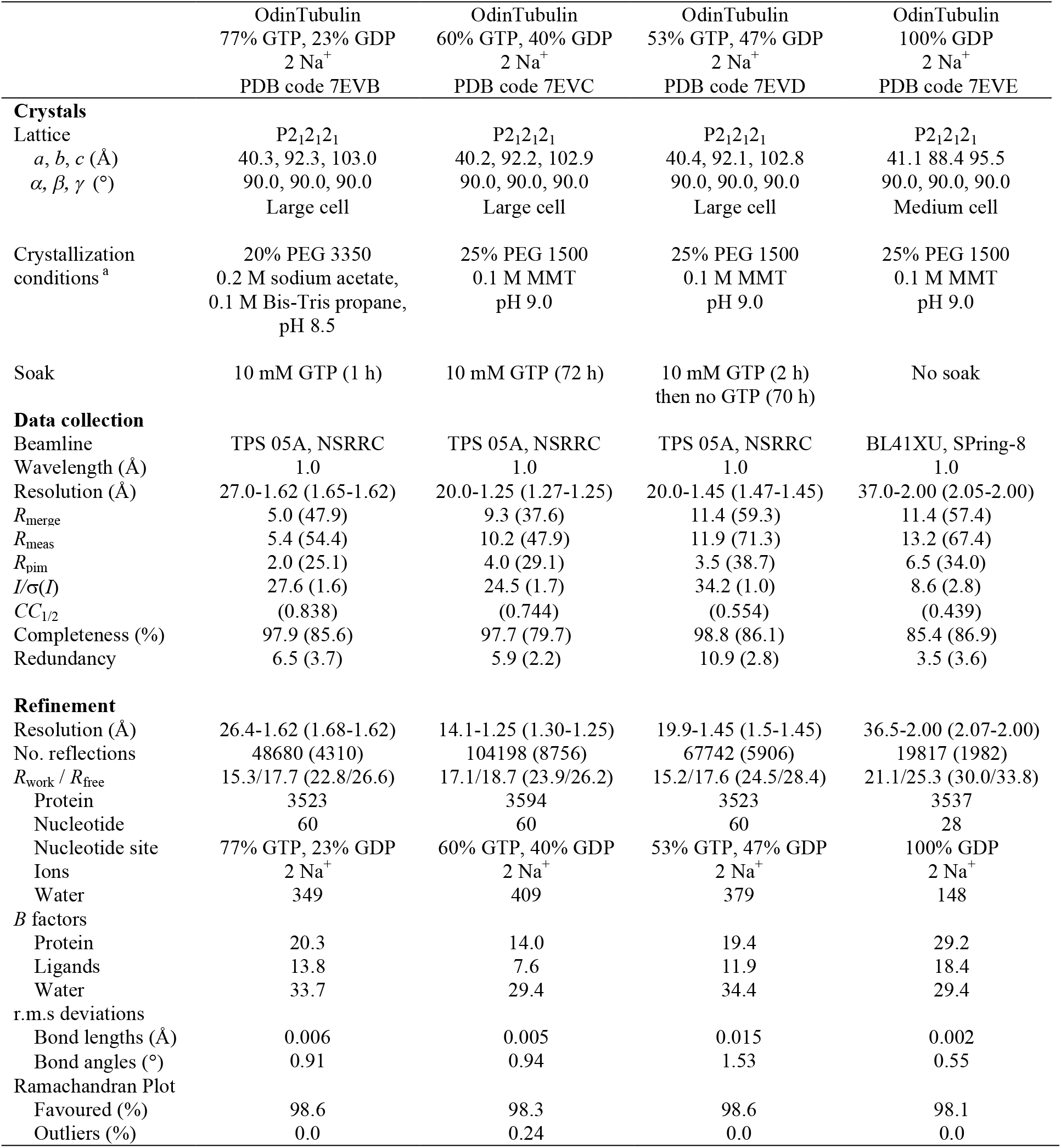
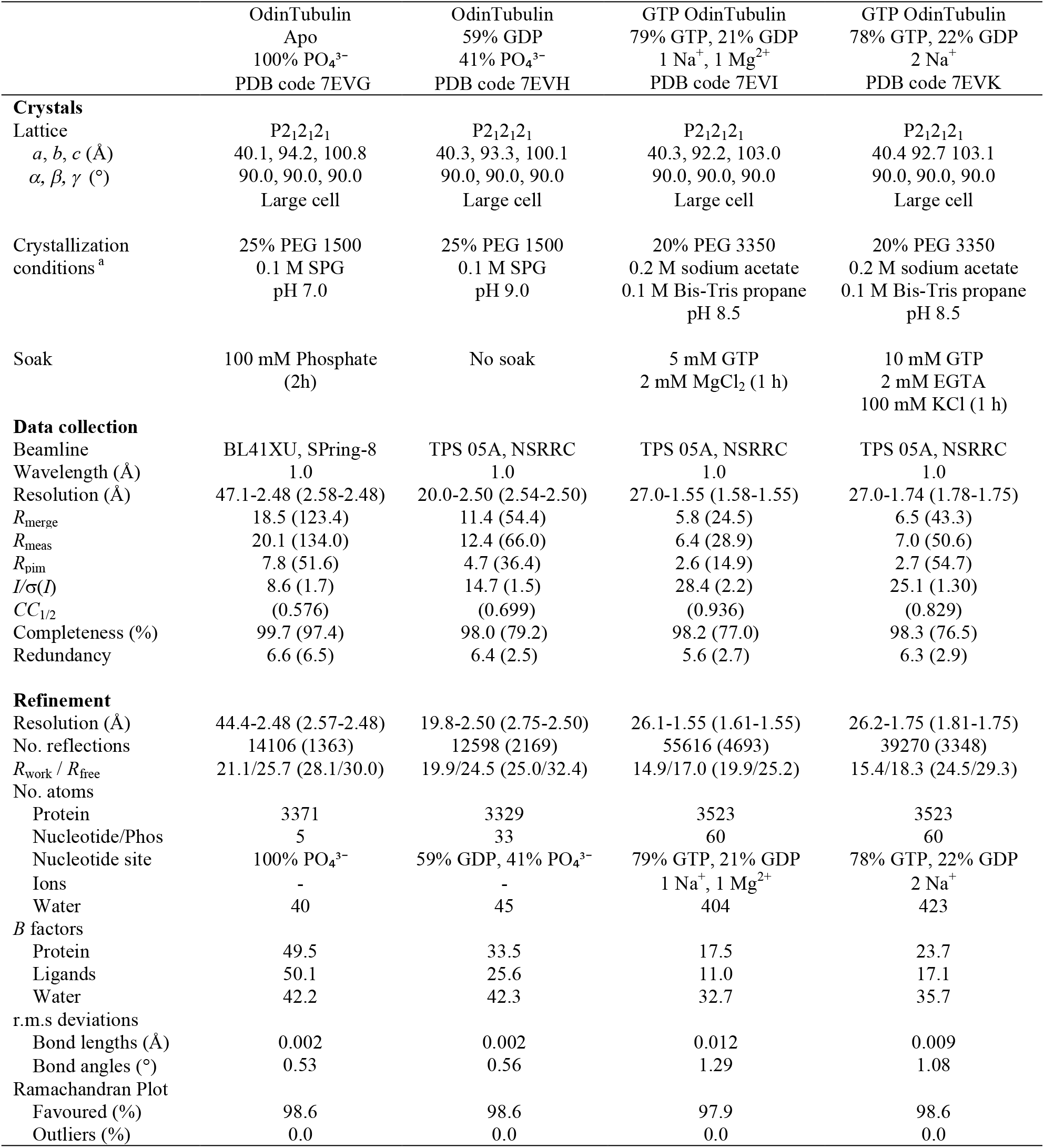
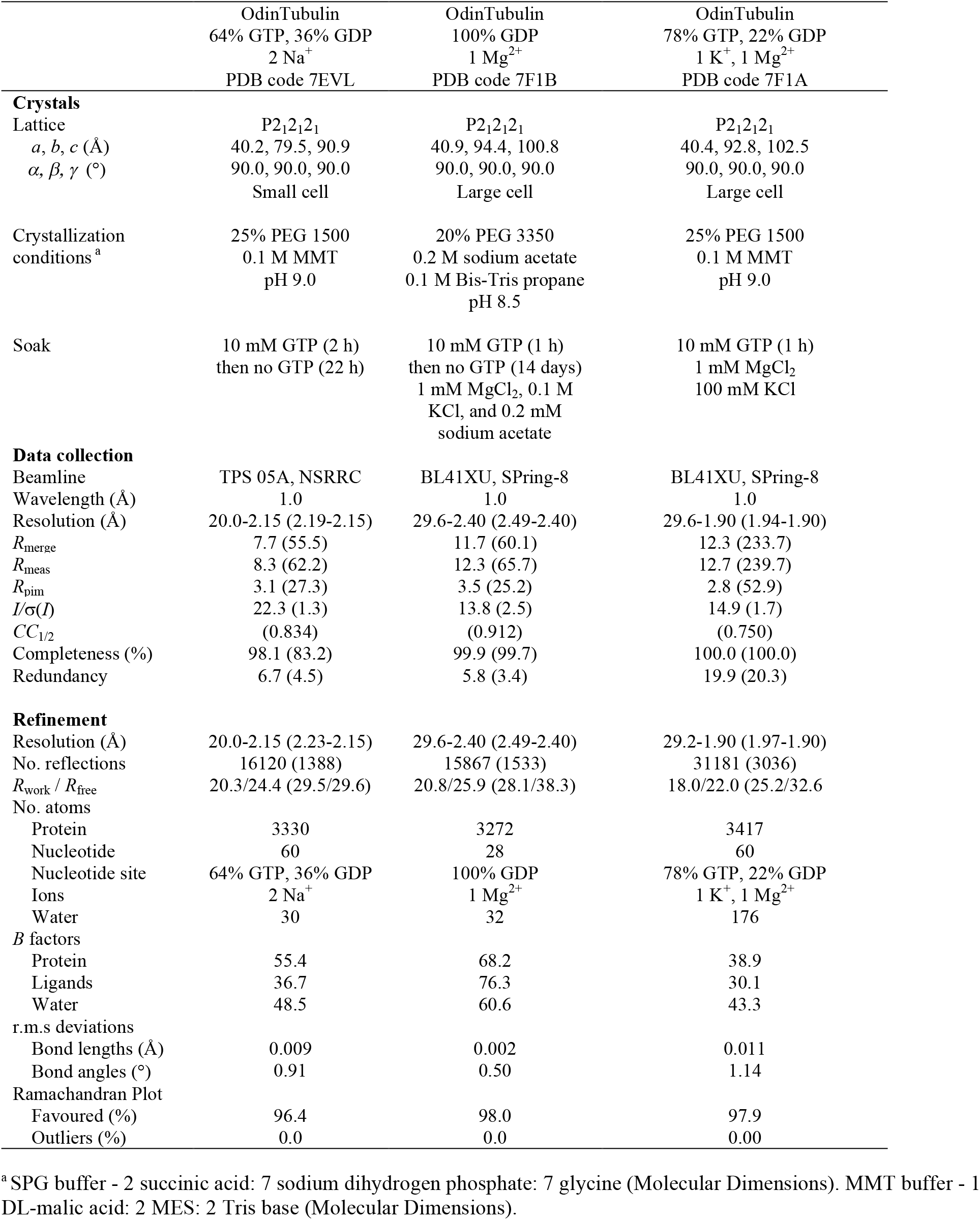
Crystallization and X-ray data collection and refinement statistics.

**Table S2.** Structural similarity of the GTP-bound OdinTubulin protomer to eukaryotic tubulins and prokaryotic FtsZs and CetZs.

|  |  |  |  |  |  | GTP-bound OdinTubulin |  |  |
| --- | --- | --- | --- | --- | --- | --- | --- | --- |
| Structure | PDB | Subunit | Nucleotide | Resolution/Å | Method | RMSD/Å | Residues | Z-Score |
| *Eukaryotic Tubulins* |  |  |  |  |  |  |  |  |
| Stu2p-bound $\alpha/\beta$ tubulin | 4u3j-A | $\alpha$ | GTP | 2.8 | X-ray | 2.0 | 410 | 52.4 |
| tubulin-stathmin-TTL | 4iij-A | $\alpha$ | GTP | 2.6 | X-ray | 2.1 | 407 | 50.0 |
| Stu2p-bound $\alpha/\beta$ tubulin | 4u3j-B | $\beta$ | GTP | 2.8 | X-ray | 1.8 | 403 | 54.2 |
| tubulin-stathmin-TTL | 4iij-B | $\beta$ | GDP | 2.6 | X-ray | 1.9 | 406 | 54.2 |
| Gamma tubulin | 3cb2-A | $\gamma$ | GDP | 2.3 | X-ray | 2.2 | 407 | 49.8 |
| *Eukaryotic Microtubules* |  |  |  |  |  |  |  |  |
| Deacetylated microtubule | 6o2r-A | $\alpha$ | GTP | 3.3 | EM | 1.5 | 407 | 54.3 |
| Microtubule | 6dpu-A | $\alpha$ | GTP | 3.1 | EM | 1.5 | 407 | 54.0 |
| Deacetylated microtubule | 6o2r-B | $\beta$ | GDP | 3.3 | EM | 1.4 | 410 | 57.0 |
| Microtubule | 6dpu-B | $\beta$ | GMPCPP | 3.1 | EM | 1.4 | 410 | 56.5 |
| *Prokaryotic FtsZ/CetZ/TubZ* |  |  |  |  |  |  |  |  |
| CetZ *M. thermophila* | 3zid-B | CetZ | GDP | 2.0 | X-ray | 2.5 | 309 | 32.1 |
| CetZ *H. volcanii* | 4b45-A | CetZ2 | GTPγS | 2.1 | X-ray | 2.4 | 308 | 32.7 |
| CetZ *H. volcanii* | 4b46-A | CetZ1 | GDP | 1.9 | X-ray | 2.6 | 309 | 32.8 |
| FtsZ *M. jannaschii* | 1w5a-A | FtsZ | GTP | 2.4 | X-ray | 3.3 | 298 | 26.4 |
| FtsZ *S. aureus* | 3vo8-A | FtsZ | GDP | 2.25 | X-ray | 3.2 | 279 | 23.1 |
| FtsZ *M. tuberculosis* | 1rlu-A | FtsZ | GTPγS | 2.1 | X-ray | 2.8 | 288 | 26.9 |
| TubZ *B. thuringiensis* | 2xka-A | TubZ | GTPγS | 3.0 | X-ray | 3.7 | 315 | 24.5 |

Matching numbers of residues, RMSD and Z-Score indicate the structural similarity to OdinTubulin. The most similar structures are highlighted in blue.

**Table S3.** Structural similarity of a pair of GTP-bound OdinTubulin protomer to pairs of eukaryotic tubulins.

| Structure | PDB | Subunit | Nucleotide | Resolution/Å | Method | RMSD/Å | Residues |
| --- | --- | --- | --- | --- | --- | --- | --- |
| *Eukaryotic Tubulins* |  |  |  |  |  |  |  |
| Stu2p-bound $\alpha/\beta$ tubulin | 4u3j-A | $\alpha\beta$ | GTP | 2.8 | X-ray | 2.3 | 749 |
| tubulin-stathmin-TTL | 4iij-A | $\alpha\beta$ | GTP | 2.6 | X-ray | 2.4 | 766 |
| tubulin-stathmin-TTL | 4iij-B | $\beta\alpha$ | GDP | 2.6 | X-ray | 2.6 | 752 |
| *Eukaryotic Microtubules* |  |  |  |  |  |  |  |
| Deacetylated microtubule | 6o2r | $\alpha\beta$ | GTP | 3.3 | EM | 1.3 | 799 |
| Microtubule | 6dpu | $\alpha\beta$ | GTP/GTP | 3.1 | EM | 1.6 | 800 |
| Deacetylated microtubule | 6o2r | $\beta\alpha$ | GDP/GTP | 3.3 | EM | 1.3 | 800 |
| Microtubule | 6dpu | $\beta\alpha$ | GMPCPP | 3.1 | EM | 1.8 | 786 |

Matching numbers of residues and RMSD values indicate the structural similarity. The most similar structures are highlighted in blue.

**Movie S1. Comparison of the GTP-bound OdinTubulin protofilament with the GDP-bound microtubule**. Superimposition, based on the lower protomer, of the two GTP-bound OdinTubulin (7EVB) symmetry-related subunits from the crystal packing (yellow) onto two subunits of eukaryotic tubulin (cyan) from the GDP-bound microtubule (PDB 6o2r). A complimentary video to Fig. 1C and fig. S3.

**Movie S2. Comparison of the GTP-bound OdinTubulin protofilament with the GMPPCP-bound microtubule**. Superimposition, based on the lower protomer, of the two GTP-bound OdinTubulin (7EVB) symmetry-related subunits from the crystal packing (yellow) onto two subunits of eukaryotic tubulin (cyan) from the guanosine-5’-[(α, β)-methyleno]triphosphate (GMPPCP)-bound microtubule (PDB 6dpu). A complimentary video to Fig. 1C and fig. S3.

**Movie S3. Comparison of the GTP-bound OdinTubulin protofilament with the stathmin-bound curved protofilament**. Superimposition, based on the lower protomer, of the two GTP-bound OdinTubulin (7EVB) symmetry-related subunits from the crystal packing (yellow) onto two subunits of eukaryotic tubulin cyan) from the stathmin-bound curved protofilament (PDB 4iij). A complimentary video to Fig. 1C and fig. S3.

**Movie S4. The OdinTubulin subunit interactions in the protofilament**. A complimentary video to Fig. 2A. Two subunits of GTP-bound OdinTubulin (7EVB) are depicted. The α7 helix and preceding loop (blue) and α8 helix and preceding loop (red) comprise the nucleotide sensor motif, which connects the upper and lower GTP-binding sites (sticks). Secondary structure elements are colored by domain: N-terminal (pink), intermediate (cyan), and C-terminal (orange).

**Movie S5. The OdinTubulin interactions around GTP (7EVB) and GDP (7EVE) in the protofilament. GTP 7EVB**, A complimentary video to Fig. 2C. Green, black and cyan spheres indicate a magnesium ion, a sodium ion and water molecules, respectively. The proposed hydrolytic water is shown as a red sphere, and the potential hydrogen receiving water molecule in blue. The purple dashed line indicates the direction for nucleophilic attack on the GTP γ-phosphate. The OMIT map is shown contoured at 1 σ around the structure (black), with the proposed hydrolytic water density highlighted in cyan. **GDP constrained 7F1B**, A complimentary video to Fig. 2D. In the GDP-bound structure within the protofilament packing, three water molecules (purple) replace the GTP γ-phosphate. The OMIT map (grey) and 2Fo-Fc map (lime green) are shown contoured at 1 σ. The OMIT map around several of the water molecules and cations is relatively weak relative to the mainchain and GDP electron density, suggesting partial occupancy. The OMIT map shows no density for one of the water molecules (purple) and very weak density in the 2Fo-Fc map at this contour level. This water is within bonding distance of the cations and is included in the structure to highlight the partial occupancy of the cation coordination.

**Movie S6. The OdinTubulin subunit interactions in the pseudo protofilament**. A complimentary video to Fig. 2F. Two subunits of phosphate bound OdinTubulin (7EVG) are depicted. The α7 helix and preceding loop (blue) and α8 helix and preceding loop (red) comprise the nucleotide sensor motif, which connects the upper and lower GTP-binding sites (sticks). Secondary structure elements are colored by domain: N-terminal (pink), intermediate (cyan), and C-terminal (orange).

**Movie S7. The conformational changes in OdinTubulin**. A complimentary video to Fig. 2G. The video shows a morph between the GTP-bound state (7EVB) and the alternate apo state (7EVG) of OdinTubulin. The α7 helix and preceding loop (blue) and α8 helix and preceding loop (red) comprise the nucleotide sensor motif, which connects the upper and lower GTP-binding sites (sticks). The domain domains are coloured N-terminal (pink), intermediate (cyan), and C-terminal (orange). The GTP molecules are depicted from the GTP-bound state for reference. F222 (upper), N226 (upper), D249 (lower) and E251(lower) are shown as yellow sticks, which highlights the mechanism of nucleotide sensing between the upper and lower nucleotide-binding sites.

**Movie S8. Polymerization of OdinTubulin followed by IRM**. Wide field view of the elongation of microtubules and OdinTubulin under the conditions outlined in Fig. 5.

**Movie S9. Polymerization of OdinTubulin followed by IRM**. A zoomed view of the elongation of microtubules and OdinTubulin under the conditions outlined in Fig. 5.

